# Comparing Heritability Estimators under Alternative Structures of Linkage Disequilibrium

**DOI:** 10.1101/2021.09.08.459523

**Authors:** Alan Min, Elizabeth Thompson, Saonli Basu

## Abstract

SNP heritability of a trait is the proportion of its variance explained by the additive effects of the genome-wide single nucleotide polymorphisms (SNPs). The existing approaches to estimate SNP heritability can be broadly classified into two categories. One set of approaches model the SNP effects as fixed effects and the other treats the SNP effects as random effects. These methods make certain assumptions about the dependency among individuals (familial relationship) as well as the dependency among markers (linkage disequilibrium, LD) to provide consistent estimates of SNP heritability as the number of individuals increases. While various approaches have been proposed to account for such dependencies, it remains unclear which estimates reported in the literature are more robust against various model mis-specifications. Here we investigate the impact of different structures of LD and familial relatedness on heritability estimation. We show that the performance of different methods for heritability estimation depends heavily on the structure of the underlying pattern of LD and the degree of relatedness among sampled individuals. However, contrary to the claim in the current literature, we did not find significant differences in the performance of these fixed-SNP-effects and random-SNP-effects approaches. Moreover, we established the equivalence between the two method-of-moments estimators, one from each of these two lines of approaches.

## 1 Introduction

Fundamental to the study of inheritance is the partitioning of the total phenotypic variation into genetic and environmental components Visscher et al. (2008). Using family studies, the phenotypic variance-covariance matrix can be parameterized to include the variance of an additive genetic effect, and an environmental effect (Lynch et al., 1998). Specific family designs, such as twin studies can accommodate both shared and nonshared environmental effects. The ratio of the genetic variance component to the total phenotypic variance is the proportion of genetically controlled variation and is termed as the ‘narrow-sense heritability’. As shown in the recent review of more than 17,000 twin studies (Polderman et al., 2015), heritability provides useful information on the power to identify causal genetic markers in a genome-wide association study (GWAS), is used to estimate familial recurrence risk of disease, and informs the genetic architecture of the trait (e.g., through partitioning by genomic region or tissue-specific expression).

GWASs seek to understand the relationship between these traits and millions of single-nucleotide polymorphisms (SNPs), a type of genetic variant. Linear models are widely used in the field of statistical genetics to assess both individual and cumulative contribution of genetic variants on a trait. The individual contribution is assessed by treating each variant as a fixed effect (fixed-SNP-effect model) while adjusting for relevant covariates in a linear regression (Dicker, 2014; Schwartzman et al., 2019; Bulik-Sullivan et al., 2015) or by treating each variant as a random effect (random-SNP-effect model) by using a linear mixed effect model (Yang et al., 2010, 2011; Speed et al., 2012). The first line of approaches treat the sample of seemingly unrelated people as independent, but the random-SNP-effect models attempt to estimate the genetic relatedness among individuals to improve the efficiency of estimation of genetic variance. Nowadays, with the increasing ability to sequence many genetic variants in large cohort studies (UK Biobank Bycroft et al. (2018), Precision Medicine cohort Collins & Varmus (2015), and the Million Veterans Program Gaziano et al. (2016) are a few such examples), there is significant interest to estimate the cumulative contribution of the genome-wide causal variants. Often we assess such cumulative contribution by estimating the proportion of variance explained by the additive effects of the causal variants in the genome; that is, the “SNP heritability”. Estimating this SNP heritability provides an upper bound to the variance explained by genetic prediction models. However, regardless of the choice to model individuals as independent or not, all approaches make certain assumptions to model the cumulative genetic contributions, which are often violated in practice. This paper takes a closer look at the impact of violations of assumptions on the properties of estimators.

The random-SNP-effect models assume an infinitesimal model for the SNP effects and use of genome-wide SNP data on distantly related individuals (Yang et al., 2010, 2011; Lee et al., 2011; Yang et al., 2012; Lee et al., 2012; Speed et al., 2012; Bulik-Sullivan et al., 2015; Haseman & Elston, 1972) to estimate the pairwise genetic relatedness between sampled individuals. These approaches assume that each causal SNP makes a random contribution to the phenotype, and these contributions are correlated between individuals who have similar genotypes. By partitioning the phenotypic covariance matrix among all individuals into a genetic similarity matrix and a random variation matrix, the approach estimates the proportional contribution of the genetics to the total phenotypic variation. The estimation of the heritability parameter heavily depends on the estimation of a high-dimensional genetic relationship matrix (GRM). The matrix is usually estimated from the observed data on *M* markers for all *n* individuals in the cohort. Two methods of estimation are used to estimate heritability under this model. One is a likelihood-based approach, which includes GCTA (Yang et al., 2010) and LDAK (Speed et al., 2012). The other approach uses method-of-moments technique to estimate heritability, such as HE regression (Haseman & Elston, 1972). A major advantage of this mixed effect model approach is that it can account for related individuals, but the general recommendation is to exclude individuals with relatedness greater than 0.025 in the estimation of heritability (Yang et al., 2011).

The other set of approaches assume SNP effects are arbitrary and fixed (Dicker, 2014; Schwartzman et al., 2019). The proposed estimators are consistent and asymptotically normal in high-dimensional linear models with Gaussian predictors and errors, where the number of causal predictors *m* is proportional to the number of observations *n*; in fact, consistency holds even in settings where *m/n* → *ρ*, where *ρ* is an element of (0, ∞). These set of approaches cannot easily accommodate relatives in the model, and thus the consistency of the estimator is derived under the assumption that the sampled individuals have independent genotypes.

Since the true genetic architecture of any given trait is unknown, existing methods are susceptible to bias. In this paper, we show that differences in such genetic architecture can yield vastly different estimates of SNP-heritability for a trait, even when applied to the same data. There are methods that allow fitting multiple variance components to accommodate the genome wide linkage disequilibrium (LD) and minor allele frequencies (MAF) differences, but it is not obvious if stratifying by MAF/LD produces more accurate estimates of total SNP-heritability. Moreover, the likelihood-based approaches become computationally intensive on large Biobank level datasets. On the other hand, the fixed-effect model based approaches can explicitly model MAF- and LD-dependent architectures when estimating SNP-heritability; however, these approaches can produce drastically different estimates when their assumptions are violated (Speed et al., 2012; Gusev et al., 2013; Ma & Dicker, 2019). Thus, it remains unclear which estimates of genome-wide SNP-heritability computed from Biobank-scale data (e.g., UK Biobank) are more reliable.

The choices of fixed- and random-effects modeling of genetic effects have been discussed several times in the context of heritability estimation (Gibson, 2012). It has been argued that the fixed-SNP-effect model approaches can handle the linkage disequilibrium (LD) among the markers in a more statistically rigorous way, whereas the random-SNP-effect models correct for LD in an adhoc manner (Ma & Dicker, 2019). This paper derived a link between the random-effects model methods and the fixed-effect approaches through the use of the Mahalanobis kernel.

A thorough study of the effects of LD and the impact of including related individuals could provide us a better understanding of the specificity and sensitivity of different approaches to heritability estimation. In this paper, we present a robust set of simulation studies with varying structures of LD and compare the performance of a wide array of estimators. We further provide some theoretical results that justify the observed simulation performance. We demonstrate through theoretical derivations as well as simulation studies that the potential impact of LD on a random-effect-model based estimator (Haseman & Elston, 1972) depends on the extent and structure of correlation of the causal and non-causal variants. We also show that a version of the fixed effect model estimator proposed by Dicker (2014) is essentially equivalent to the Haseman-Elston method-of-moments estimator (Haseman & Elston, 1972). Our findings in this paper did not demonstrate any particular advantage of the fixed-SNP-effect models over the random-SNP-effect models in presence of LD, at least for the case where heritability is estimated using only common variants.

This rest of the paper is organized as follows: first, different methods to estimate heritability are explained and analytical formulae to compare their performance under different LD structures are presented. We then describe strategies to simulate LD and relatedness structure and to evaluate both fixed and random effects models. Finally, results are presented and discussed.

## 2 Methods

### 2.1 Genotypes, Phenotypes, and Heritability Estimation

We consider a set of *n* individuals from a homogeneous population, typed at *M* SNP markers, assumed to be in Hardy-Weinberg equilibrium. Note that notation is also listed Table 1. Assume an n × M matrix of genotypes **G** = (*G*_*ij*_), where *G*_*ij*_ = 0, 1, 2 is the number of copies of the reference allele for individual *i* at locus *j* with population frequency *p*_*j*_. Thus *G*_*ij*_, *i* = 1, 2, …, *n*, has mean 2*p*_*j*_ and variance 2*p*_*j*_(1 − *p*_*j*_), *j* = 1, 2, …, *M*. The vector of standardized genotypes for individual *i* at marker *j* is given by

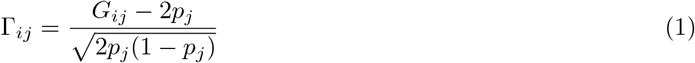

so that Γ_*ij*_ has mean 0 and variance 1.

**Table 1:** Glossary of notation used

| Variable | Definition |
| --- | --- |
| $n$ | Number of individuals in an analysis |
| $M$ | Total number of markers in an analysis |
| $m$ | The number of causal markers in an analysis. A marker is considered causal if it has a non-zero direct effect on phenotype. |
| $i, k$ | Typically used to index individuals, $i, k = 1, \dots, n$. |
| $j, \ell$ | Typically used to index markers, $j, \ell = 1, \dots, m$ or $M$. |
| $p_j$ | The population frequency of locus $j$ |
| $\sigma_g^2$ | The variance in phenotype attributable to genotypic effects |
| $\sigma_e^2$ | The variance in phenotype attributable to environment. |
| $r_{jl}$ | The genotypic correlation between loci $j$ and $l$ in the population |
| $\phi_{ik}$ | Genotypic correlation between individuals $i$ and $k$ |
| $\mathbf{G}$ | A matrix of genotypes with $n$ rows and $M$ columns |
| $\mathbf{\Gamma}_A$ | A matrix of normalized genotypes with $n$ rows and $M$ columns (Equation (1)) |
| $\mathbf{\Gamma}_C$ | A matrix formed by the $m$ columns of $\mathbf{\Gamma}_A$ that correspond to the causal markers |
| $\mathbf{\Psi}$ | The GRM calculated using all markers; an $n \times n$ matrix. $M^{-1} \mathbf{\Gamma}_A \mathbf{\Gamma}_A'$. |
| $\mathbf{\Sigma}$ | The $M \times M$ marker LD matrix calculated from all markers. $n^{-1} \mathbf{\Gamma}_A' \mathbf{\Gamma}_A$ |
| $\boldsymbol{\beta}$ | An $m$-vector of effects of causal loci on phenotype, or sometimes an $M$-vector augmented by 0's (Equation (4)) |
| $\mathbf{y}$ | An $n$-vector of phenotypic values of individuals. |

The matrix of standardized genotypes for all markers, **Γ**_*A*_ = (Γ_*ij*_), carries information on the relatedness of individuals, and the LD among markers. While E(Γ_*ij*_Γ_*iℓ*_) = *r*_*jℓ*_ is the genotypic correlation between loci within an individual, E(Γ_*ij*_Γ_*kj*_) = *ϕ*_*ik*_ measures the genotypic correlation between individuals. We define the GRM **Ψ** as in Yang et al. (2010)

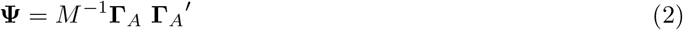

and we define the LD matrix as

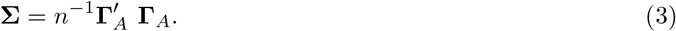

where we use the single quote (′) to denote the transpose of a matrix. In large samples, the empirical allele frequencies 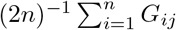 can be used as an estimate of the population frequency *p*_*j*_ of Equation (1) in forming the matrices **Ψ** and **Σ**.

Suppose that the first *m* of the *M* markers are *causal*, having a direct impact on phenotype. We denote by **Γ**_*C*_ the matrix consisting of the *m* columns of **Γ**_*A*_ corresponding to the causal markers, and adopt the classical trait model of Fisher (1918). The phenotype of individual *i* is given by

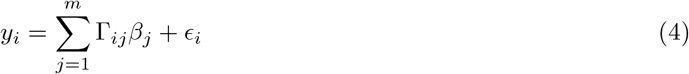

where *β*_*j*_, Γ_*ij*_ and *ϵ*_*i*_ are mutually independent and have mean 0. Then E(*y*_*i*_|**Γ**_*C*_) ≡ 0 so that var(y_i_) = E(var(y_i_ | **Γ**_C_)). In SNP heritability estimation, 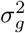 is the phenotypic variance attributable to SNPs, and 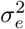 is the phenotypic variance attributable to environmental factors. In a random-SNP-effect model, we assume 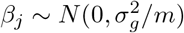, and 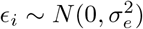. Then

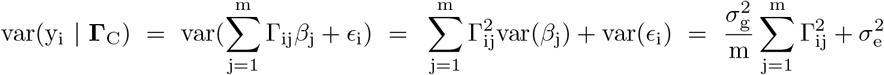

Then either conditionally on **Γ**_*C*_ as *m* becomes large, or taking expectations over Γ_*ij*_, 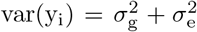. On the other hand, a fixed-SNP-effect model assumes that ***β*** is a fixed quantity, with *β*_*j*_ = 0 for non-causal markers. In that case, we define 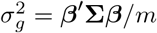, and note E(Γ_ij_Γ_i*ℓ*_) = Σ_j*ℓ*_ so that

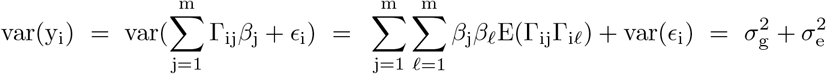

Thus in either case, the phenotypic variance 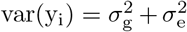 and SNP-heritability is 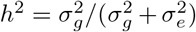. If phenotypes are standardized to have variance 1, then 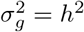 and 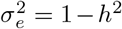. More generally, estimation of heritability is primarily concerned with estimation of 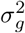, the estimate of *h*^2^ being then obtained by dividing by the empirical variance of the phenotypes *y*_*i*_, *i* = 1, …, *n*.

### 2.2 Overview of Estimators

In our overview of the methodologies for heritability estimation, we concentrate on method-of moments estimation and likelihood-based estimation for the random-SNP-effect models. We further compare these estimators with the fixed-SNP-effect model based estimators (Dicker, 2014; Schwartzman et al., 2019). The Supplementary Material provides more details on these estimators.

For the likelihood methods, we consider the GCTA (Yang et al., 2011) and LDAK (Speed et al., 2012) approaches. In brief, GCTA is a random-SNP-effect model derived under asumptions similar to those of Section 2.1. The approach uses REML (Patterson & Thompson, 1971) to estimate 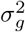 and 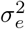. It estimates heritability assuming that phenotypes are drawn from a multivariate Normal distribution, where the log likelihood function is

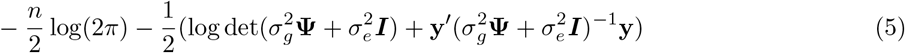

LDAK (Speed et al., 2012) uses a similar model, except reweighting the SNP markers to adjust for LD. More details on the GCTA and LDAK approaches can be found in Supplementary Section S1.

For the method-of-moments estimators, we first considered a Haseman-Elston (HE) estimator (Haseman & Elston, 1972; Yang et al., 2011), an estimator from the random-effect approach category. The estimator of 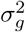, derived in the Supplementary Section S2.1, has the form

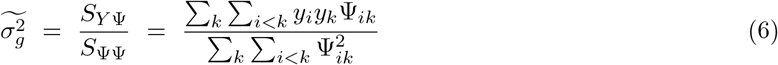

An estimate of heritability is then given by dividing by the empirical variance of phenotypes *Y*_*i*_. Further properties of this estimator in the case of no LD are given in Supplementary Section 2.1.

We also considered two method-of moments estimators from the fixed-effect approach category, Dicker-1 and Dicker-2 (Dicker, 2014). Dicker-1 is applicable in the case of no LD. It is derived and discussed in the Supplementary Section 2.2, and takes the form

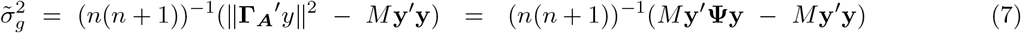

We consider this estimator primarily for comparison with the HE estimator: see Supplementary Section S2.1 and Section 2.3.2.

In presence of LD, if **Σ** is invertible, the estimator of Dicker (2014) becomes

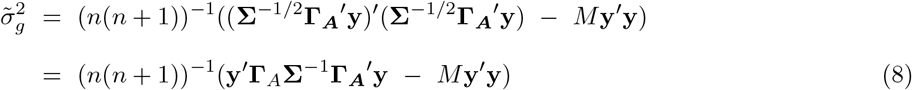

However, in many cases, **Σ** is not invertible because *M > n*. In these cases, to address the LD, Dicker (2014) derives an estimator which we denote Dicker-2. This estimator uses moments of the trace of the LD matrix **Σ** to correct for LD, resulting in an estimator of 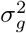

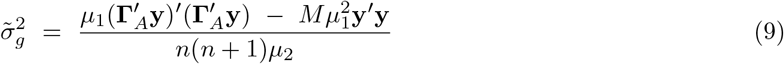

where

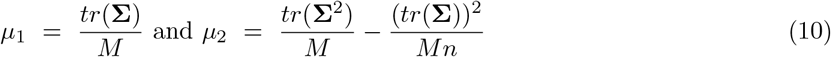

Further details of the Dicker-2 estimator are given Supplementary Section S3.2.

### 2.3 Impact of Linkage Disequilibrium

#### 2.3.1 Impact of LD on the Haseman-Elston Estimator

In this section we consider the impact of marker mispecification and marker LD on the numerator and denominator of the HE estimator, and hence on the estimate of 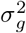. We assume unrelated individuals but correlated markers, so that E(Γ_ij_Γ_k*ℓ*_) = 0 if *i* ≠ *k*, but E(Γ_ij_Γ_i*ℓ*_) = r_j*ℓ*_, with −1 ≤ *r*_*jℓ*_ ≤ 1, and *r*_*jj*_ = 1.

We split the markers into *m* causal markers *C* and (*M* − *m*) non-causal markers *F*. Note that all markers are used in the GRM: Ψ = *M* ^−1^**Γ**_*A*_ **Γ**′_*A*_, but that only causal markers **Γ**_*C*_ contribute to the phenotype **y**. For convenience, assume that the first *m* markers are causal: *C* = {1, …, *m*} and *F* = {(*m* + 1), …, *M*}. Then, following the same derivation as in Supplementary Section S2.1, for *i* ≠ *k* we obtain,

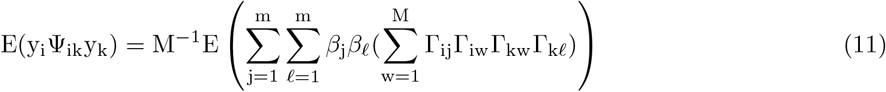

If the *β*_*j*_ have mean 0, and are uncorrelated, we have only terms in *j* = *ℓ*, and this reduces to

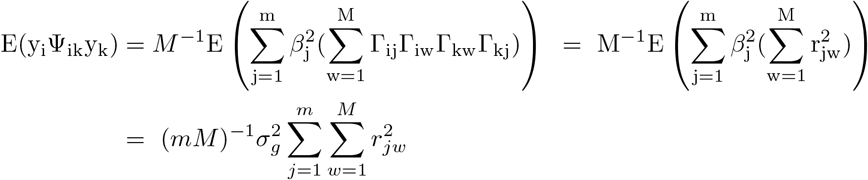

here using that individuals *i* and *k* are independent, and that 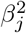 has expectation 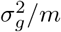.Then

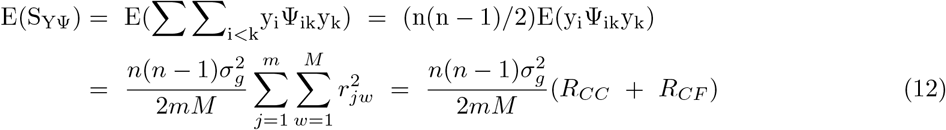

where for convenience we denote the sums of squared correlations

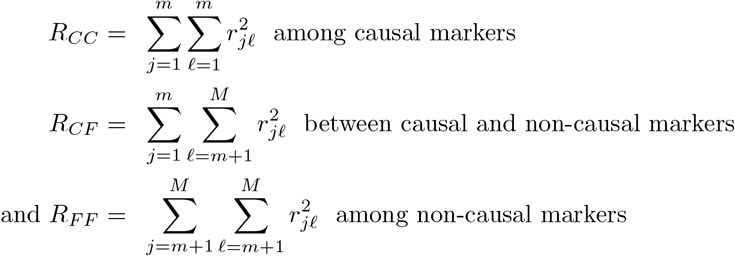

Considering similarly the denominator of the HE estimator,

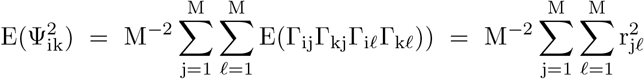

so that

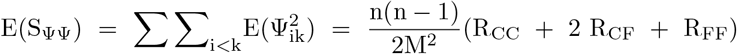

leading finally to the ratio of expectations of *S*_*Y* Ψ_ and *S*_ΨΨ_

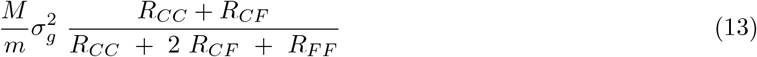

Equation (13) covers many cases of interest. First, if there is no LD, *R*_*CC*_ = *m, R*_*CF*_ = 0, and *R*_*FF*_ = (*M* − *m*), giving the results of Section S2. Second, if the GRM contains only causal markers *M* = *m*, then LD among these causal markers does not cause bias. Third, if additional markers *F* are not in LD with each other, nor with the causal markers *C, R*_*CF*_ = *R*_*FF*_ = 0, and again no bias results. Note that generally inclusion of additional markers in the GRM is less serious than omission of causal markers. If **Γ**_*A*_ is missing causal markers *j* then Equation (11) will not include the contributions of those *β*_*j*_ and *S*_*Y* Ψ_ will be decreased, but *S*_ΨΨ_ will not (on average) be affected, leading to underestimation of 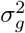.

In other cases, typically there is bias. If causal markers are in regions of high LD, then (per marker) *R*_*CC*_ dominates over *R*_*FF*_, and 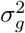 will be overestimated, while if causal markers are in regions of low LD *R*_*FF*_ in the denominator will dominate, and 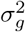 will be underestimated. In some special cases biases cancel out. Consider first a special case of causal markers in regions of “average LD”; suppose all *r*_*jℓ*_ = *s* for *j* ≠ *ℓ*. Then *R*_*CC*_ = *m* + *m*(*m* − 1)*s*^2^, *R*_*CF*_ = *m*(*M* − *m*)*s*^2^ and *R*_*FF*_ = (*M* − *m*) + (*M* − *m*)(*M* − *m* − 1)*s*^2^ and some arithmetic shows there is no bias. Two other examples occur in the simulations of Section 3. In both the autocorrelation and block simulations, causal and non-causal markers are alternating. Then *M* = 2*m* and *R*_*FF*_ = *R*_*CC*_, and Equation (13) again shows there is no bias. This is demonstrated in the simulation results in Figures 1 and 2. While these exact situations are unlikely to occur in reality, these examples show that some forms of LD may not have a huge impact on the bias of the HE estimator.

**Figure 1:**
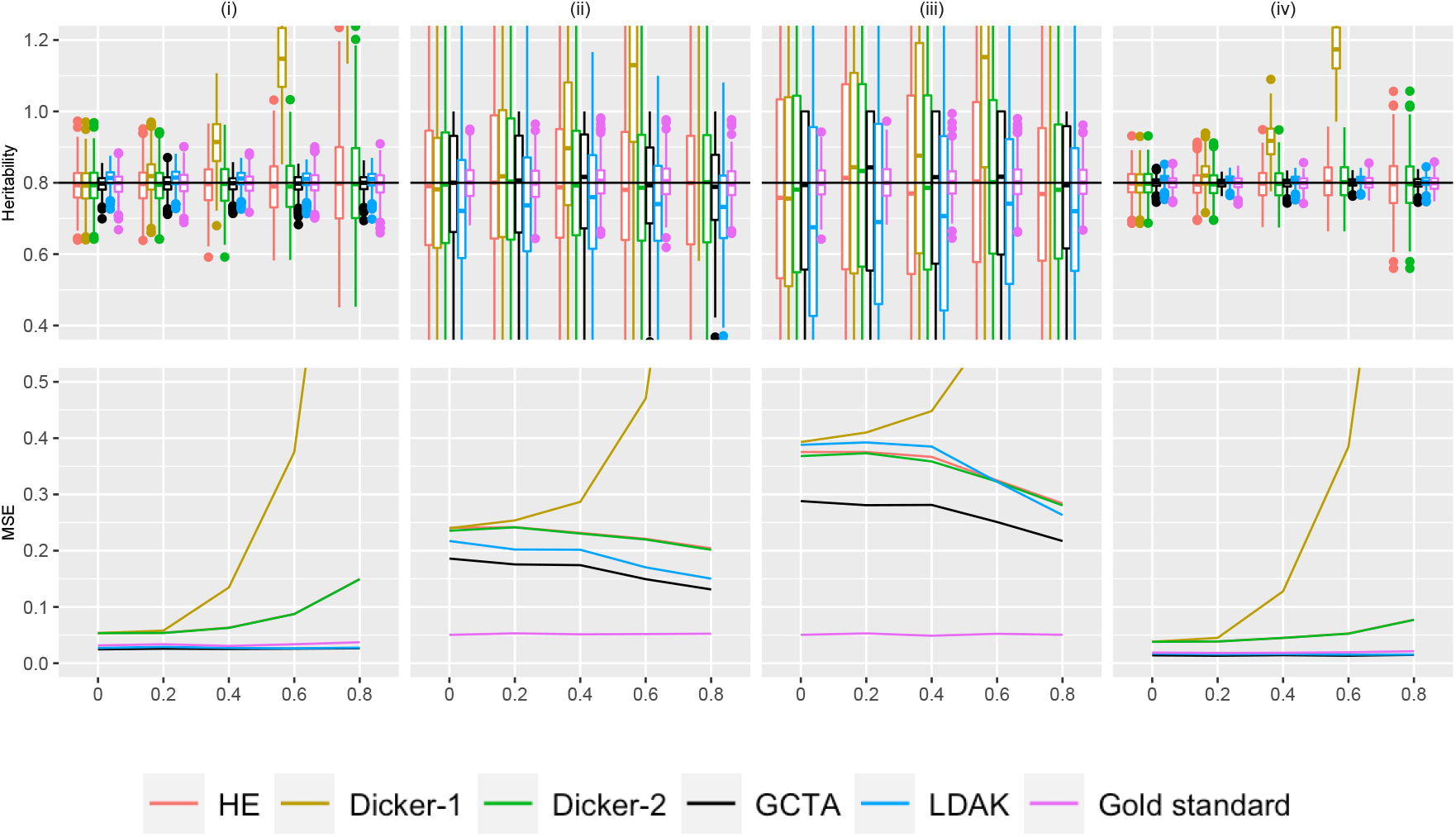
Simulation Study 1A (autocorrelated markers). On the top row, the X-axis plots the parameter *ρ*, the autocorrelation correlation coefficient between simulated markers as described in Supplementary Section S4. Estimates of *h*^2^ using different estimators are plotted along the Y-axis. The value *n* refers to the number of individuals simulated. The value *M* is the total number of markers simulated, where half of the markers are causal, set in an alternating fashion, as described in Section 2.5. We consider (i) *n* = 1000, *m* = 100 (ii) *n* = 200, *m* = 500, (iii) *n* = 200, *m* = 1500 (iv) *n* = 2000, *m* = 500. 500 data sets were simulated for each condition. A horizontal line is shown at *h*^2^ = .8, the simulated truth. On the bottom row, the X-axis is the parameter *ρ*, and the MSE of each of the estimators is plotted on the Y-axis.

**Figure 2:**
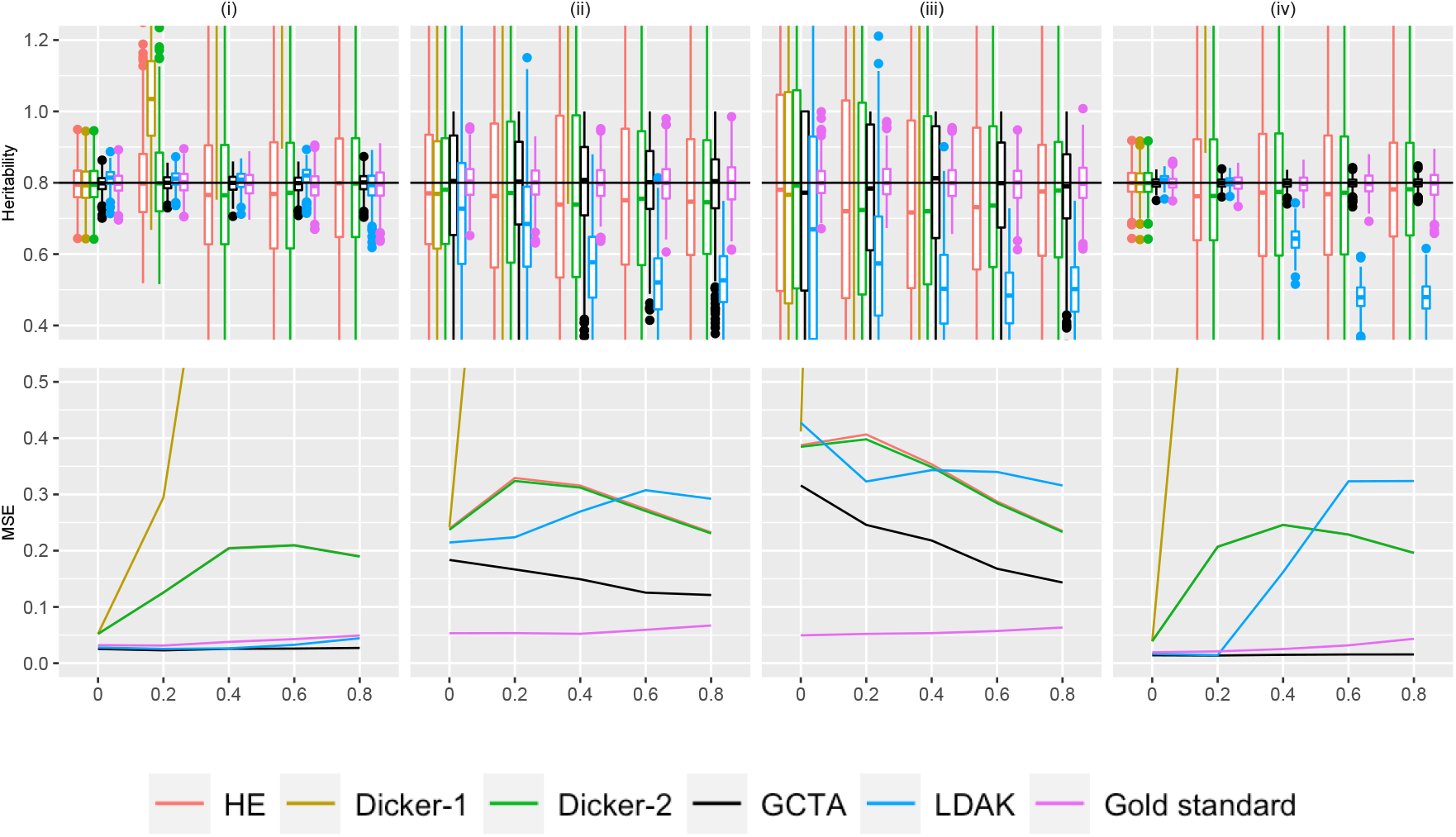
Simulation Study 1B (block markers). On the top row, the X-axis plots the parameter *ρ*, the block correlation coefficient between simulated markers as described in Supplementary Section S4. Estimates of *h*^2^ using different estimators are plotted along the Y-axis. The value *n* refers to the number of individuals simulated. The value *M* is the total number of markers simulated, where half of the markers are causal, set in an alternating fashion, as described in Section 2.5. We consider (i) *n* = 1000, *m* = 100 (ii) *n* = 200, *m* = 500, (iii) *n* = 200, *m* = 1500 (iv) *n* = 2000, *m* = 500. 500 data sets were simulated for each condition. A horizontal line is shown at *h*^2^ = .8, the simulated truth. On the bottom row, the X-axis is the parameter *ρ*, and the MSE of each of the estimators is plotted on the Y-axis.

The case of duplication of markers also considered in the simulations is different, and Equation (13) again provides an estimate of the bias. In this example, there is no LD in the *m* causal markers, so *R*_*CC*_ = *m*. The genotypes at subset of *d* of these markers are replicated *r* additional times, but these replicates are non-causal. So *M* = *m* + *rd. R*_*CF*_ = *rd* and *R*_*FF*_ = *r*^2^*d*. Then Equation (13) reduces to

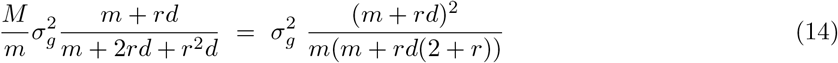

Note that if no markers are replicated (*d* = 0) or all markers are replicated (*d* = *m*) then there is no bias. Note also that the result only depends on the proportion of markers replicated. If *d* = *gm* then Equation (14) reduces to 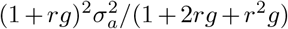. Although the expectation of the ratio is approximated by the ratio of expectations in Equation (13), simulation shows this approximation gives an accurate estimate of the bias: see Supplementary Section 3.1 (Figure S2).

#### 2.3.2 Impact of LD on the Dicker-1 Estimator

We consider now the estimator of Equation (7) in the presence of LD. Even in the absence of LD, this estimator of 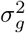 is unbiased only if 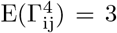 (Supplementary Section S2.2) but the bias is negligible for large *M* or *n*. Note that *E*(Γ_*ij*_Γ_*il*_) = Σ_*jl*_ so that

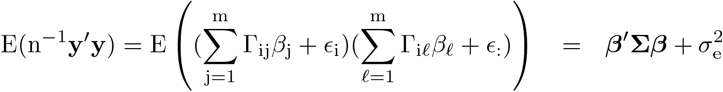

where the vector ***β*** is augmented by zeros for the non-causal markers, *j* = (*m* + 1), …, *M*. The fixed-effects Dicker-1 estimator thus estimates *τ* ^2^ = ***β***′**Σ*β*** which may differ from the additive genetic variance 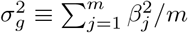 in the presence of LD. Additionally

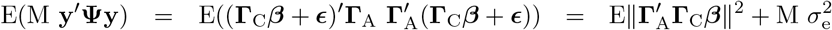

which is not easily considered in terms of the fixed-effects model.

However, in our simulation studies, each replicate uses *β*_*j*_ at causal loci *j* = 1, …, *m* that are independently generated 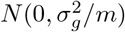 and independent of the standardized genotypes Γ_*ij*_ (Section 2.5). Thus, over replicate simulations

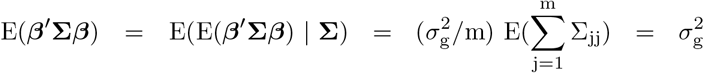

and we may consider expectations of the estimator (7) over replicates under the assumption that *β*_*j*_ (*j* = 1, …, *m*) are independent 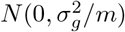. Then 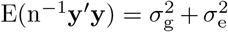 and 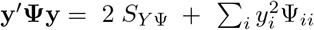 so that, from Equation (12)

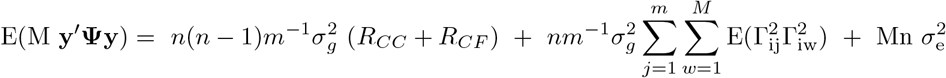

In general 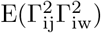 is unknown, but a lower bound on the double-sum term is the no-LD value *mK* +*m*(*M* − 1) (see Supplementary Section S2.2) while a rough approximation might be *mK* + (*M* − 1)(*R*_*CC*_ + *R*_*CF*_), where again 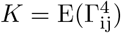. This approximation gives the overall result

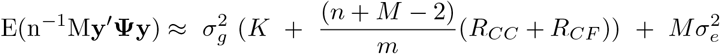

and the approximate expectation of the estimator (7) as

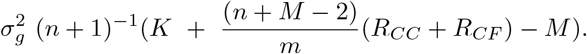

Since the squared correlations 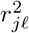 are non-negative, (*R*_*CC*_ + *R*_*CF*_) ≥ *m* and the estimator will overestimate 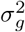 and hence also heritability *h*^2^. Unlike the HE estimator where the LD inflates both numerator and denominator (Equation (13), the form of the estimator (7) means that it can only be inflated by LD.

#### 2.3.3 Equivalence of Haseman-Elston and Dicker-2

As will be shown in Section 3.2, estimates from the Dicker-2 and HE regression were very similar, although Dicker-2 explicitly models LD. Analytically, under certain normalization schemes, the two estimators are effectively equivalent. This suggests that efforts to correct for LD in the Dicker-2 framework do not ensure improved performance of this estimator compared to the HE estimator.

We begin by reformulating HE regression. We recall from Equation (6) that the HE estimator is given by

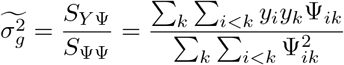

We can rewrite this in matrix form, giving us

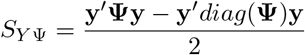

Under Hardy-Weinberg Equilibrium, the GRM should have values that are approximately 1 on the diagonal. We assume *y* is normalized to have variance 1, which results in

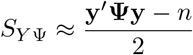

Now we consider *S*_ΨΨ_. Noting that 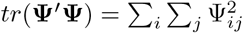,

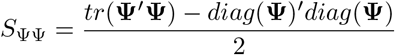

Using the same approximation on the GRM, our equation is

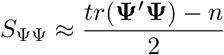

Together, the HE estimator is approximately

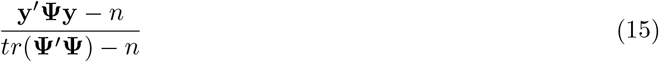

Now consider the equations for Dicker 2. First note that in Equations (9) and (10), *µ*_1_ ≈ 1 since the genotypes have variance 1. Next, we use the cyclic property of traces that *tr*(*ABCD*) = *tr*(*DABC*) for matrices *A, B, C* and *D*

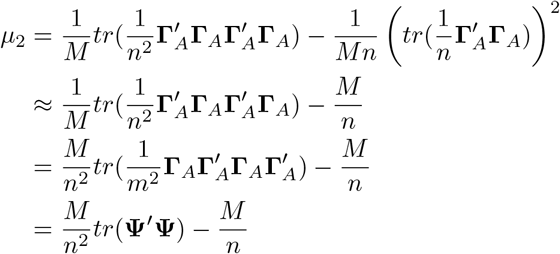

Since *n* ≈ *n* + 1, for large *n*, we have that the Dicker 2 estimator (Equation 9) is approximately

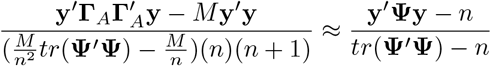

Which is the same as Equation (15).

### 2.4 Impact of Relatedness of Individuals on Moments Estimators

Under the assumption of independence of individuals, the SD of the estimator of 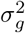 or of *h*^2^ increases with the number of markers *M* (Supplementary Section S2). This arises because in the limit, the matrix **Γ**_*A*_ converges in probability to the identity matrix, rendering *h*^2^ non-identifiable. However, this is an artefact of the assumption of complete independence (unrelatedness) of individuals. In any real sample, regardless of the extent of correction for population structure, there will always be variation in the degree of relatedness of individuals, even if any single pairwise relatedness measure is small. Note that the original formulation of HE estimators (Haseman & Elston, 1972) made use of the genetic similarity between known relatives. In this section, we therefore consider the case where individuals may be related, so standardised genotypes Γ_*ij*_ and Γ_*kj*_ are no longer independent. For simplicity we ignore LD: that is Γ_*ij*_ and Γ_*kℓ*_ are independent, whether or not *i* = *k*.

Under relatedness and inbreeding it remains the case that E(Γ_ij_) = 0, but var(Γ_ij_) = (1 + F_i_) and E(Γ_ij_Γ_kj_) = *ϕ*_ik_, where *F*_*i*_ is the inbreeding coefficient of individual *i*, and *ϕ*_*ik*_ is the relatedness of *i* and *k*, or twice the coefficient of kinship between *i* and *k*. To consider the HE estimator (6), for *i* ≠ *k*,

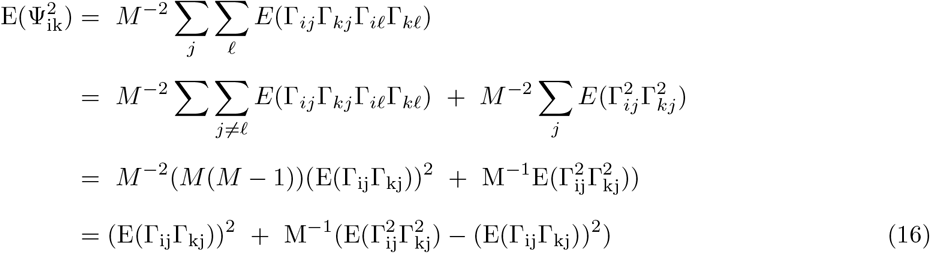

Hence as *M* → ∞ *S*_ΨΨ_ tends to

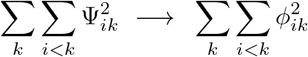

and *S*_*Y* Ψ_ tends to

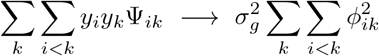

Thus, contrary to the results of Supplementary Section S2.1 for unrelated individuals, the SD of the HE estimator no longer increases as *M* → ∞, but rather will depend on the magnitude of 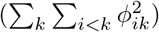. Although this sum may be small, if even any of the *ϕ*_*ik*_ are non-zero it is strictly positive, and eventually relatedness will bound the SD of the estimator of 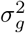.

Relatedness poses greater problems for the Dicker-1 estimator (Equation 7) which involves the diagonal terms of the GRM matrix Ψ. Considering the expected quadratic form

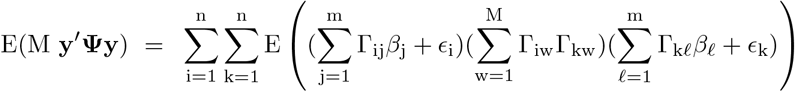

Now, even the coefficient of 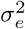 is no longer *mn* but 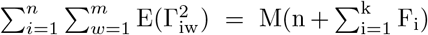 while that of 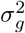 is, as in Supplementary Section 2.2

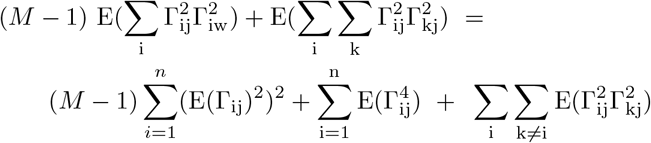

This expectation now involves not only (1 + *F*_*i*_)^2^, and 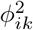 but also higher order moments.

Although the derivation of distributional properties of the the Dicker method-of-moments estimators depends critically on the assumption of 2*n* independent genomes, there is nothing in the derivation of Section 2.3.3 that assumes **Ψ** is diagonal. Indeed, the trace equation

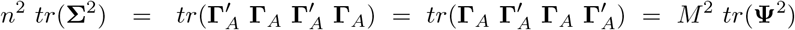

used in showing the approximate equivalence of the HE and Dicker-2 estimators, suggests that the Dicker-2 accommodation of LD in the absence of relatedness is alternatively accommodating relatedness in the absence of LD. Thus, as will be seen in the results of Section 3.4, the close equivalence of the Dicker-2 and HE estimators should hold under relatedness, and, as seen from equation (16) above, the standard deviation will no longer increase indefinitely as *M* → ∞.

### 2.5 Simulation strategy

We performed simulation studies to assess the impact of LD structure and relatedness of individuals on heritability estimation. Each simulated data set consisted of genotypes **G** at *M* markers (*m* causal markers) for *n* unrelated individuals. The marker allele frequencies were those of a randomly chosen subset of markers from the 1000 genomes project from Chromosome 1 in the AFR population (Clarke et al., 2017). This set of frequencies was filtered to have allele frequency less than .95 and greater than .05 and was fixed over data set simulations.

Genotypes are standardized using their empirical allele frequencies. Phenotypes were simulated for *n* individuals, given their genotypes at the *m* causal markers, in accordance with the linear model of Equation (4):

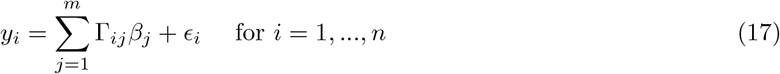

For the chosen value of *h*^2^, (0 < *h*^2^ < 1), the *m*-vector of genetic effects ***β*** was simulated with independent components *β*_*j*_ ∼ *N* (0, *h*^2^*/m*) for *j* = 1, …, *m*. The independent residual effects *ϵ*_*i*_ ∼ *N* (0, 1 − *h*^2^) for *i* = 1, …, *n*. Thus, for the purposes of the simulation 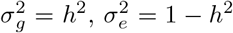, and var(*y*_*i*_) = 1, with *h*^2^ set to 0.8 for all simulations. (see Section 2.1).

For each simulated dataset, we implemented the Dicker, and Haseman Elston estimators in R Version 4.0.2 as described in the Supplementary Section S2. We used GCTA Yang et al. (2011) and LDAK Speed et al. (2012) as representative likelihood estimators, both of which are described in more detail in Supplementary Section S1. We also report a gold standard estimator to assess the performance of these different methods. The gold standard estimate is calculated assuming we know the true values of ***β***: the empirical variance of **Γ**_***C***_***β*** is divided by the empirical variance of the phenotypes. This gold standard estimator can be expressed as

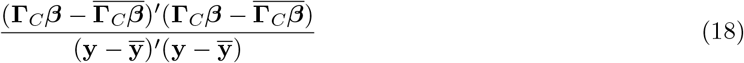

In <u>simulation study 1</u>, we assessed the impact of different LD structures on heritability estimation. We generated genotypes assuming three kinds of LD structure: autocorrelated, block, and repeat. More details of the LD structures are given in Supplementary Section S4. Each data set was simulated with a new ***β*** and **G**. For each LD structure, we studied the impact of both sample size *n* and the number of causal markers *m* on heritability estimation, and simulated data sets at five levels of LD. For each LD structure and level, we generated 500 simulated data sets.

For the autocorrelation and block structures, we considered the following combinations of *n* and *m*: (1) *n* = 1000, *m* = 100, (2) *n* = 200, *m* = 500, (3) *n* = 200, *m* = 1500, and (4) *n* = 2000, *m* = 500. Comparing (1) and (2) provides insight on differences in estimates of *h*^2^ depending on if *n > m* or *m > n*, whereas (2) and (3) compares estimates with different number of causal markers, and (2) and (4) compares estimates with different numbers of individuals. We first generated genotypes at *M* = 2*m* markers. We used marker correalations *ρ* = 0, 0.2, 0.4, 0.6, and 0.8., as detailed in Supplementary Section S4 (Note that *ρ* = 0 is the no-LD case.) The markers were then assigned to be alternating causal and non-causal (*m* = *M/*2).

For the repeat structure, we considered the cases: (1) *n* = 1000, *m* = 200, (2) *n* = 200, *m* = 1000, (3) *n* = 200, *m* = 3000, and (4) *n* = 2000, *m* = 1000. In this case, we first simulated genotypes for the *m* independent causal markers. The genotypes at the first 10% of markers were then repeated *r* times, where *r* = 0, 2, 4, 6, or 8. (Note that *r* = 0 is the no-LD case.) The repeat copies of the markers are non-causal, so the number of non-causal markers is 0.1*rm*, and *M* = *m* + 0.1*rm*. In Supplementary Figure S3 (panels C & F) the first *m* markers are causal, and the last (*M* − *m*) are the non-causal repeat copies.

In <u>simulation study 2</u>, we investigated the behavior of likelihood models by plotting log-likelihood values (Equation 5) as a function of 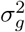 and 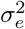. The GRM **Ψ** in Equation (5) was calculated using Equation (2). Of interest was the relationship between the shape of the log-likelihood function and the number of individuals and causal markers, and the shape of the likelihood as the number of repeats increased. From the results of simulation study 1, we hypothesized that the shape would be different when *m > n*, where GCTA underestimated heritability, compared to when *m < n*, where GCTA overestimated (comparing (i) and (iv) of Figure 3). The combinations of numbers of markers and individuals was the same as with the repeats in Simulation Study 1, and the allele frequencies were taken from the AFR sample of the 1000 Genomes Project, as before. We include plots with no repeated markers (Figure 4A) and with 10% of the markers repeated 8 times (Figure 4B).

**Figure 3:**
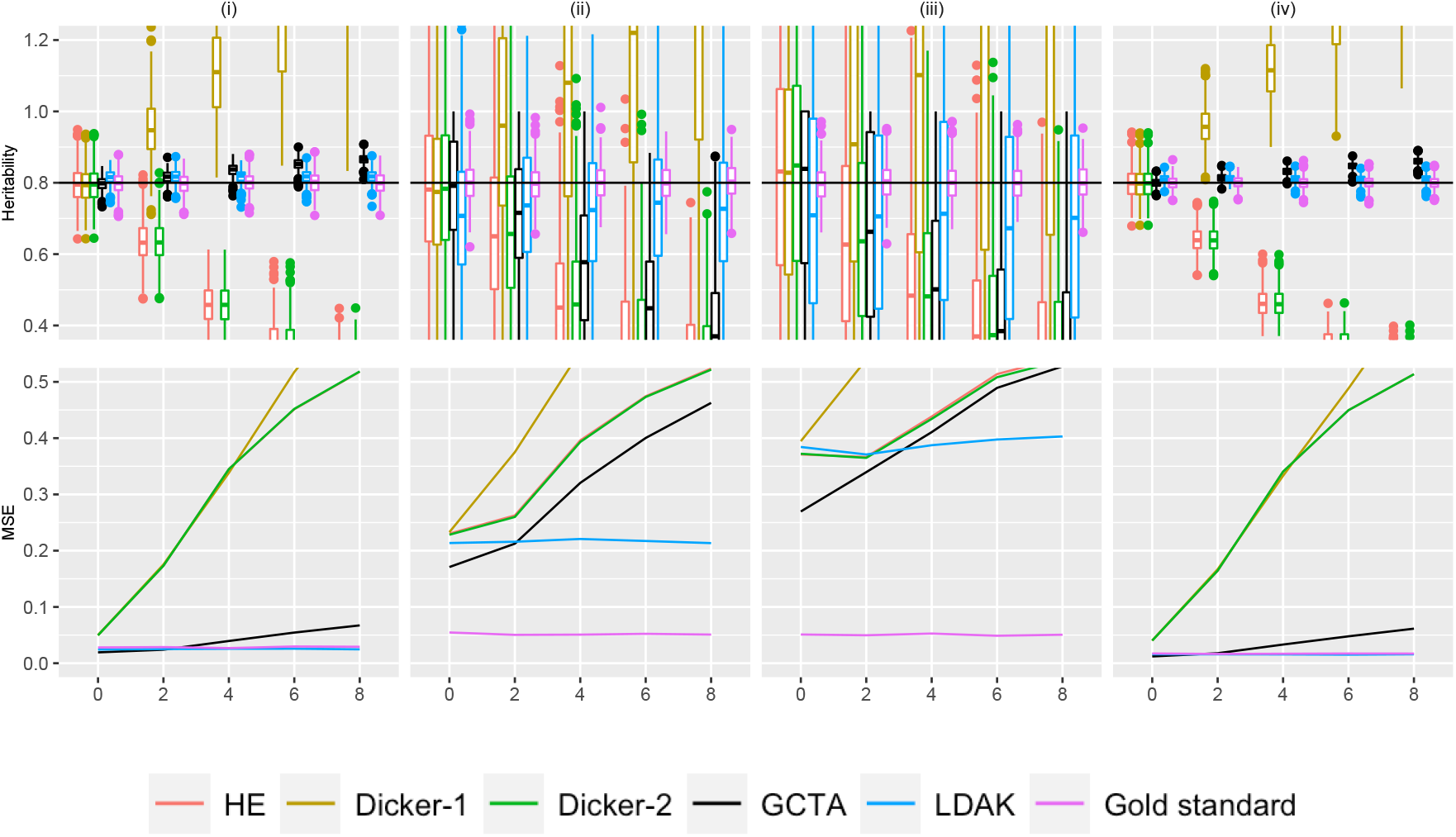
Simulation Study 1C (repeated markers). On the top row, the X-axis plots the parameter *r*, the number of times that 10% of the markers are being repeated as described in Supplementary Section S4. Estimates of *h*^2^ using different estimators are plotted along the Y-axis. The value *n* refers to the number of individuals simulated. The value *m* is the total number of causal markers simulated, as described in Section 2.5. We consider (i) *n* = 1000, *m* = 200 (ii) *n* = 200, *m* = 1000, (iii) *n* = 200, *m* = 3000 (iv) *n* = 2000, *m* = 1000. 500 data sets were simulated for each condition. A horizontal line is shown at *h*^2^ = .8, the simulated truth. On the bottom row, the X-axis is the parameter *r*, and the MSE of each of the estimators is plotted on the Y-axis.

**Figure 4:**
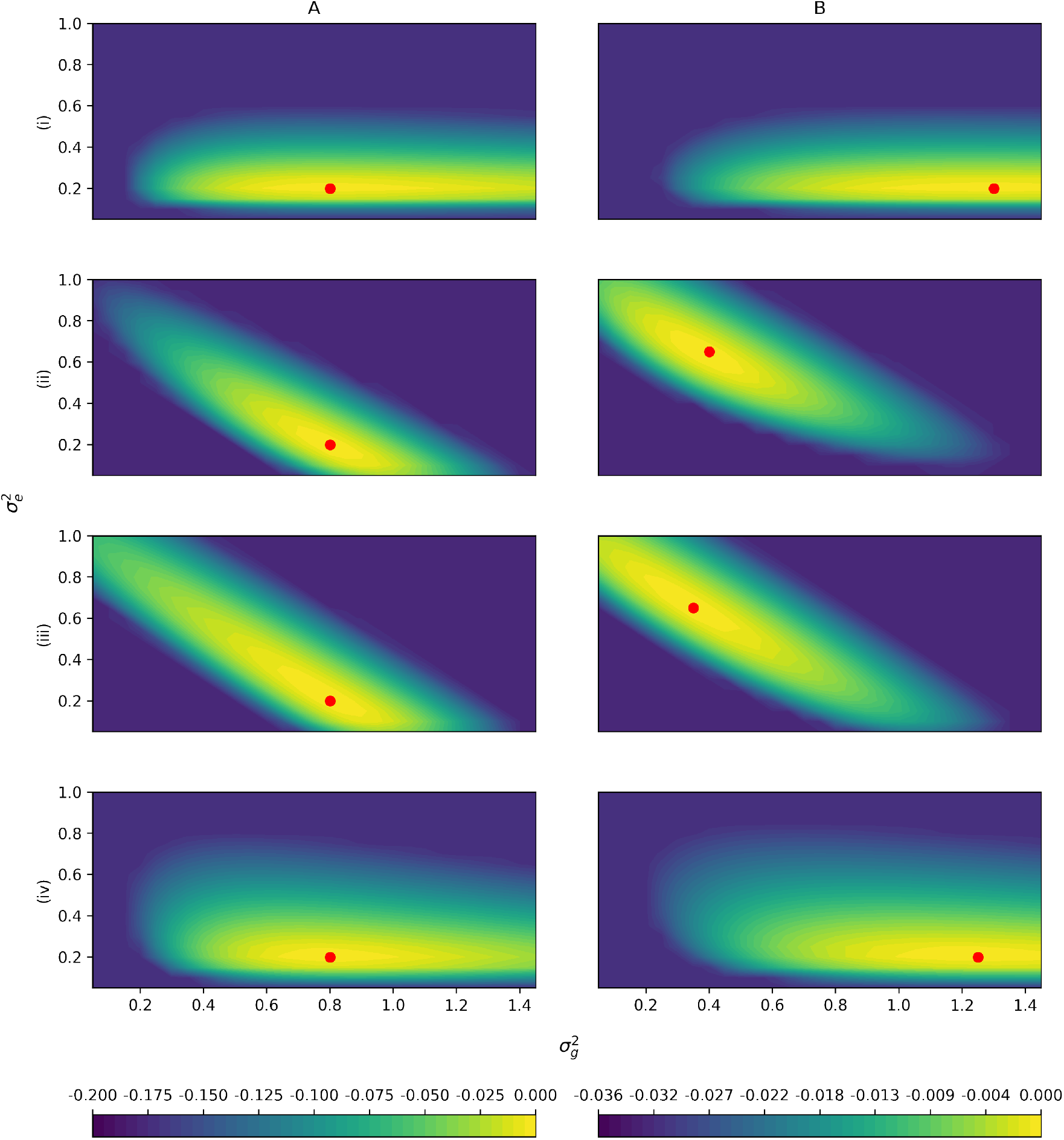
Simulation Study 2. The difference of the log likelihood from the maximum log likelihood is plotted for parameters 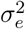 on the Y-axis and 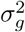 on the X-axis. The colors depict the value of the difference from the maximum log likelihood. Likelihoods are truncated at the 60% quantile of B(i) for rows (i) and (iv), and at the 60% quantile of A(ii) for rows (ii) and (iii) for visibility. Row labels correspond with Figure 3, with (i) *n* = 1000, *m* = 200 (ii) *n* = 200, *m* = 1000, (iii) *n* = 200, *m* = 3000, and (iv) *n* = 2000, *m* = 1000. Column A has markers with no LD, and in column B, 10% of the markers are repeated 8 times, corresponding the the rightmost points in Figure 3. The average of 100 independent simulations using a grid with spacing 0.05 is plotted in each panel. Note that there is one color scale shared between (i) and (iv) on the left, and a different color scale shared between (ii) and (iii) on the right due to different ranges. The red point indicates the location of the maximum likelihood.

The log likelihoods minus the maximum log likelihood were plotted. Likelihoods were truncated at the 60% quantile of B(i) for rows (i) and (iv), and at the 60% quantile of A(ii) for rows (ii) and (iii). These cutoffs were chosen because they were the plots that had the lowest 60% quantile. Plots were generated for (i) *n* = 1000, *m* = 200 (ii) *n* = 200, *m* = 1000, (iii) *n* = 200, *m* = 3000 (iv) *n* = 2000, *m* = 1000, in following with simulation study 1. We averaged log likelihoods of 100 simulated data sets with grid spacing 0.05. Due to differences in ranges, there is a shared color bar between (i) and (iv), and a different shared color bar between (ii) and (iii). A red dot is used to mark the location of the maximum log likelihood.

In <u>simulation study 3</u>, we assessed the impact of related individuals on heritability estimation. We simulated 1st, 2nd, and 3rd cousins using the rres package in R (Wang et al., 2017) as well as unrelated individuals to illustrate our findings in Section 2.4. The segment length option in rres was set to 3000 centimorgans. Using the same set of allele frequencies as previously, we simulated marker genotypes for 400 individuals, in 10 40-ships. A *k*-ship is defined to be a set of *k* cousins related to a certain degree. Each cousinship is unrelated with all other cousinships. The number of markers ranged from 400 to 4000 in steps of 400. Phenotypes were generated using Equation (17). For every combination of cousinships and number of markers, we simulated 500 sets of 10 40-ships. A visualization of the GRM of the dataset as shown in Supplementary Figure S4, using 1000 markers.

## 3 Results

### 3.1 Simulation Study 1: Bias and Variance when *ρ* = 0 or *r* = 0 (no LD)

The special case of no LD in Simulation Study 1 is shown in Figures 1, 2, and 3 in panels of the upper row at the left-hand point of each point. These figures verify that the estimators were generally unbiased in estimating the heritability. One exception is in LDAK, where when *n* = 200, LDAK seemed to underestimate heritability.

Although we generally observed no bias in the estimators under independent markers, we saw that the estimators had a wide range of variances. In the cases *n* = 1000, *M* = 200 and *n* = 2000, *M* = 1000 (columns (i) and (iv) in Figures 1-3), the variance of the GCTA, and LDAK estimators were lower than the variance of the moments estimators, but this difference is less pronounced in the cases where *n* = 200 (columns (ii) and (iii)), which may suggest that the number of individuals affects the likelihood based estimators more than the moments based estimators. The lower variance resulted in lower MSE for GCTA for all conditions with *ρ* = 0, but the bias in LDAK caused it to have comparable MSE to the moments estimators when *n* = 200 (Figures 1, 2, and 3).

We can also compare cases when the number of individuals is kept constant while the number of markers is increased by comparing *n* = 200, *m* = 500 in column (ii) vs *n* = 200, *m* = 1500 in column (iii). In Supplementary Section S2.1, we found that with unrelated individuals and independent markers, the standard deviation of heritability should be asymptotically proportional to 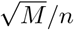 in the case of the Haseman Elston estimator. Accordingly, when the number of causal markers increases, the standard deviation of the heritability estimates increased as well. This is shown in both Figures 1, 2, and 3, where MSEs were higher in column (iii) compared to column (ii). This trend appeared to hold true for both the likelihood based estimates and the moments based estimates.

We can compare cases when the number of individuals increased while holding the number of markers constant by comparing *n* = 200, *m* = 500 in column (i) vs *n* = 2000, *m* = 500 in column (iv). The variance and MSE of the heritability estimates decreased for all estimators, which agreed with the theoretical result for the Haseman Elston estimator.

Finally, some of the biases in the behavior of LDAK may be that the LDAK model does not match our generative model. LDAK reweights their genotypes using *X*_*ij*_ = (*G*_*ij*_ − 2*f*_*j*_) × [2*f*_*j*_(1 − *f*_*j*_)]^*α*^, and *α* is recommended to be 1.25 (Speed et al., 2012, 2017). More details can be found in Supplementary Section S1. Our model doesn’t explicitly simulate phenotypes in this manner, however. To investigate this, we also chose *α* in LDAK to be −1, which matches our simulated phenotypes due to our normalization scheme (Equation 1). Results (not included) were largely similar, although the estimated heritability was slightly closer to the simulated truth in the repeat case.

### 3.2 Simulation Study 1: Impact of marker LD

#### (a) Autocorrelation Structure

Data were simulated using the autocorrelation structure as described in Section 2.5, and a representative set of moments and likelihood estimators are evaluated on these simulated data. The estimated variance and bias of different estimators is shown in Figure 1.

The HE estimator and the Dicker-1 estimator do not explicitly account for LD structure, and because the Dicker-1 estimate was developed for the no-LD case, it shows bias when LD is present. In the top row of Figure 1, the Dicker-1 estimator shown in gold showed an increase in bias as *ρ* increased for all of (i)-(iv). Consequently, the MSE of the Dicker-1 estimator increases rapidly compared to all of the other estimators as we increase *ρ* (bottom row of Figure 1). In contrast, there was no increase in the MSE of the HE estimator when markers were autocorrelated, agreeing with Section 2.3.1. In the top row of Figure 1, the estimates of *h*^2^ from the HE estimator did not appear to visually differ significantly from the true value of 0.8. This estimate behaved very similarly to the Dicker-2 estimator, despite the Dicker-2 estimator explicitly attempting to correct for LD. This is analytically shown in Section 2.3.3

The likelihood estimators in Figure 1 showed generally lower MSE and no obvious bias. The GCTA estimator is shown in black and the LDAK estimator is shown in light blue. Both of these estimators seemed to have lower MSE across all values of *ρ* than the moments estimators, as seen in the bottom row.

When *n* = 200, *m* = 3000, as *ρ* increased, there was a decrease in the MSE in all the estimators except the Dicker-1 estimator. In Figure 1, it can be seen that as *ρ* increases, the first and third quartiles of the estimates of *h*^2^ decrease. It has previously been shown that fewer causal markers leads to decreased variance (Dicker, 2014), and hence this effect may be driven by a decrease in the effective number of markers as LD increases.

#### (b) Block Structure

Figure 2 shows the estimated variance and bias in different estimators when the genotypes were simulated from the block structure with parameter *ρ*, as described in Section 2.5. Similarly to the autocorrelation structure, the Dicker-1 estimator had significant bias and high MSE, although this is expected because Dicker-1 as implemented here relates to the no LD case. The HE and Dicker-2 estimators were not as affected by the LD. In contrast to the autocorrelation, however, LDAK underestimated *h*^2^ in Figure 2, columns (ii), (iii), and (iv). In the bottom row of Figure 2, this resulted in an MSE that was comparable to that of HE and Dicker-1. GCTA estimates appeared to still largely be unbiased and produced MSEs that were lower than the other estimators. Again, it was observed that there are cases when the MSE decreases as *ρ* increases, similarly to the autocorrelation case.

#### (c) Repeat Structure

Figure 3 shows the variance and bias patterns when the genotypes were simulated from the repeat structure with parameter *r*, as described in Section 2.5. As *r* increases, the number of times that 10% of the markers were simulated increased. There were *m* causal markers simulated and *n* individuals. For example, when *m* = 1000 and *r* = 3, there were 1000 causal markers that were simulated, and the first 100 markers were repeated 3 times, leading to a total of 1300 markers that were entered into the analysis. An increased value of *r* indicates more markers that are in perfect LD with the original causal markers. We also examined behavior when repeated markers had a small probability of not being exact duplicates, and results were similar but less pronounced (results not shown).

As in Figures 1 and 2, the estimates for Dicker-1 increase rapidly as *r* increases, agreeing with analytical calculations from Section (2.3.2). In contrast to Figures 1 and 2, in Figure 3, the estimates for HE and Dicker-2 decrease as *r* increases, corresponding to Equation (14) and to results in Supplementary Figure (S2), where those equations were verified through simulation. The MSE of these two estimators also increase as *r* increases (Figure 3) and further produce very similar estimates, agreeing with analytical calculations from Section 2.3.3.

In Figure 3, the GCTA estimator produces estimates that are greater than *h*^2^ = 0.8 when *n* = 1000, *m* = 200 and when *n* = 2000, *m* = 1000, but produces estimates that are lower than 0.8 when *n* = 200, *m* = 1000, and when *n* = 200, *m* = 3000. In other words, if *n > m*, then the GCTA estimator is underestimating, and when *n < m*, the GCTA estimator is overestimating.

The LDAK estimator shows the same pattern of bias as GCTA in that as *r* increases, *h*^2^ is underestimated when *n > m* and overestimated when *n < m*. This is bias is less pronounced than with GCTA, however. In the bottom panel of Figure 3, it can be seen that as *r* increases, the MSE of LDAK appears relatively constant, whereas the MSE of GCTA is increasing when *n* = 200, *M* = 1000 or *n* = 200, *M* = 3000, as seen in columns (ii) and (iii).

### 3.3 Simulation Study 2: Likelihood Surfaces

In Figure 3, GCTA displayed an upward bias when *n > m*, and a downward bias when *n < m*. We hence hypothesized that the likelihood may appear different if *n > m* versus if *m > n*. The likelihood surface captures the joint likelihood of 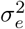 and 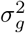. From the model in Equation (5), 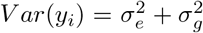. Hence we expect that the maximum likelihood lies on a diagonal, as 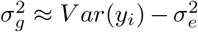. This appears to be true when the number of individuals is much larger than the number of markers, but when the number of individuals much less the number of markers, the axis of the conditional maxima becomes more horizontal (Figure 4). An intuition for this result is that as the number of individuals improves, we have better knowledge of the total phenotypic variance.

The likelihood surfaces also demonstrate a faster rate of change in the likelihood surface when the number of individuals is increased, comparing Figure 4A and G where the range of the colors is greater than in C and E. This observation corresponds with simulation study 1, where as the number of individuals increased, the variance of the estimates of heritability decreased. Finally, on the right hand side of Figure 4, the surfaces are still either diagonal or horizontal, but the maxima (red dots) are shifted. This agrees with simulation study 1 results, where there was bias in the GCTA method when the number of repeats increased.

### 3.4 Simulation Study 3: Impact of relatedness in individuals

In simulation study 3, we studied the effect of familial structure on estimates of heritability using cousinships and found that an increase in the number of causal markers generally increased MSE unless relatedness was high.

The Dicker-2, HE, and GCTA estimators appeared unbiased for each of the relatedness structures (Figure 5). For GCTA and HE, we reasoned that because their model is conditional on the GRM, it took into account relatedness. Furthermore, because we’ve shown that HE and Dicker-2 are equivalent (Section 2.3.3), we can also explain the unbiasedness of Dicker-2. LDAK was also largely unbiased in the case of unrelated individuals, but in the case of 1st cousins, as the number of markers increased, we observed that LDAK started showing downward bias.

**Figure 5:**
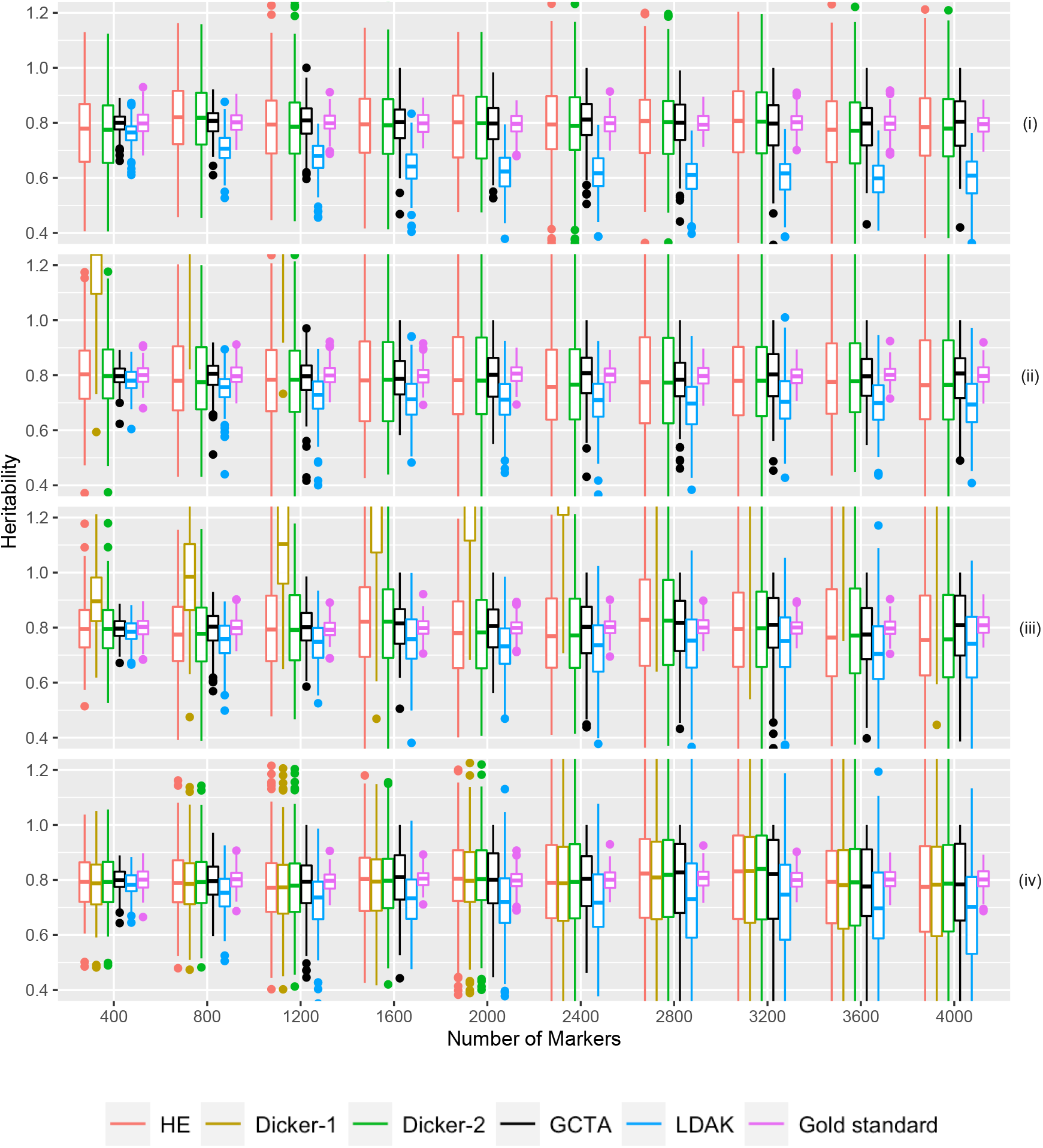
Simulation Study 3. Estimated *h*^2^ from 500 sets of 10 groups of 40 related cousins plotted on the y-axis. The number of causal markers plotted on the x-axis. Data was simulated as described in Section 3.4 Different estimators are plotted in different colors. True heritability was set to be 0.8. Note that because of the chosen range of *y* values, Dicker-1 is sometimes not visible in the figure. Panels (i), (ii), (iii), and (iv) are first-, second-, third-cousins, and unrelated individuals respectively.

For the different relatedness structures (unrelated, full sibs, first cousins) we considered, we observed similar pattern in the change of MSE as we increased the number of markers. MSE was generally the lowest when the number of markers was closer to the sample size. However as we increased the number of markers, MSE for each estimator increased (Figure 6). For HE and Dicker-2, the unrelated individuals had the lowest MSE when the number of markers was 400, but increased as the number of markers increased. On the other hand, 1st cousins had MSE that remained steady (Figure 6). When the number of markers was 4000, the MSE of the unrelated individuals was larger than the MSE of the 1st cousins, agreeing with analytical calculations from Section 2.4.

**Figure 6:**
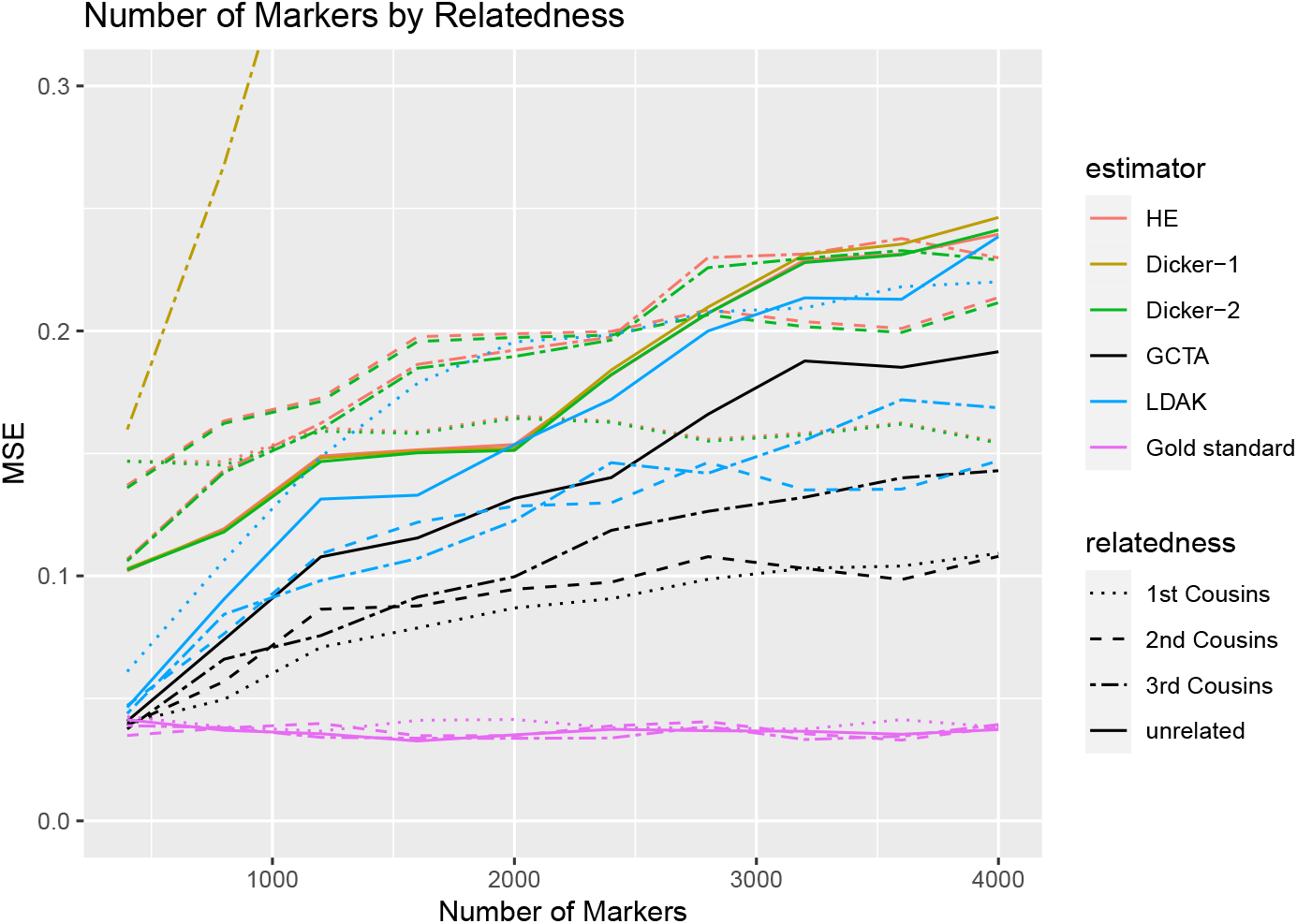
Simulation Study 3. MSE of estimates of *h*^2^ from Figure 5. The X-axis indicates the number of markers in the simulation, and the Y-axis indicates the mean square error.

For each of our estimators, unrelated individuals had the highest MSE and first cousins had the lowest MSE. Further, comparing the case of related to unrelated individuals, the MSE increased more slowly with the increase in the number of causal markers in related individuals.

## 4 Discussion

The methods for SNP heritability estimation can be broadly classified into two groups; fixed-SNP-effects models (view SNP effects as fixed effects) and the random-SNP-effects models (view the SNP effects as random effects). The fixed-SNP-effect models (Dicker, 2014; Schwartzman et al., 2019) can more easily accommodate the LD structure among the genetic variants and can accommodate variants as both causal or non-causal. However, these approaches rely on independence among the individuals in the sample. On the other hand, the random-SNP-effect models (Yang et al., 2011) can accommodate and borrow power from related individuals, though it is generally recommended to exclude relationships with higher relatedness than 0.025 (this corresponds approximately to relatives second cousins or closer) to avoid confounding due to shared environments. These random-SNP-effects models assume all variants are causal and the majority of the methods do not accommodate LD among the markers in a statistically rigorous way. The asymptotic properties of these heritability estimators depend on model assumptions. In this paper, we have studied the impact of model misspecification on heritability estimation through extensive simulation studies. We have simulated data under various LD structures and have allowed a certain portion of the variants to be non-causal. Contrary to many reports (Schwartzman et al., 2019), we found little difference in the performance of a fixed-SNP-effect model method-of-moments estimator and a MOM estimator from a random-SNP-effect model under different model misspecification.

We have derived the analytic expression for the bias of the HE estimator in presence of LD among markers. Section 2.3 considers various scenarios for the LD among causal and non-causal markers and analytically shows the impact of this correlation on the HE estimator. Our simulation studies have also considered various LD scenarios to illustrate that the bias in heritability estimation depends on the underlying LD pattern. In many scenarios, LD does not cause huge bias in the estimation. One could adopt LD-based pruning strategies to reduce the correlation among markers. However, the bias could be more severe due to omission of causal markers during LD pruning process over including these correlated markers.

In the case where Σ^−1^ can be computed (*M < n*), Dicker (2014) proposes a heritability estimator (Dicker-1) that can account for the LD among markers by rotating the genotypes. The derivation of the consistency of the estimator, however, relies on the Normality assumption. In case of large *n* and *M* (*M >> n*), our simulation studies and analytical derivation in Section 2.3.3 show that the Dicker-2 estimator (fixed-SNP-effect model based estimator) and HE estimator (random-SNP-effect model based estimator), are essentially the same. Hence, in the situation *M >> n*, Dicker-2 estimator cannot correct for the LD among markers. This is a contradiction to the claim that the Dicker (2014) always provides an improved estimator of heritability in presence of LD among markers.

Fixed-SNP-effect model based estimators generally assume that sampled individuals are independent. These approaches do not accommodate related individuals in the heritability estimation. We demonstrate that in the absence of LD, the Dicker-1 is severely inconsistent in the presence of related individuals. However, because of its equivalence to the HE estimator, the Dicker-2 estimator generates consistent estimates of heritability with related individuals in the absence of LD.

The likelihood based approaches from the random-SNP-effects model category, especially the LDAK approach showed more bias under certain model misspecification as compared to the MOM estimators. Under different LD structures, the traditional GCTA approach showed more stability in terms of both bias and precision over the LDAK estimator. We did not observe any specific advantage of adjusting for LD in the likelihood based estimators.

Under the assumption of independence of individuals, the standard errors of the heritability estimator increases with the number of causal markers. This is an artefact of the assumption of complete independence (unrelatedness) of individuals. In any real sample, regardless of the extent of correction for population structure, there will always be variation in the degree of relatedness of individuals, and the extent of variation would depend on the nature of relatedness present in the sample. As shown in Simulation 3, the precision of the heritability estimators improve if we include relatives in the sample. The MSE of the estimators were generally lower when we had certain relatedness present in the sample. Moreover, the impact of increasing the number of markers on MSE was significantly less pronounced if we had relatedness in the sample. Hence, we highly recommend to at least include second cousins, if present in the study sample, in the SNP heritability estimation. If the study sample has substantial number of first cousins, it may be beneficial to assess the sensitivity of the heritability estimate after inclusion of first cousins.

In general, MOM estimators had much larger standard errors compared to the likelihood-based estimators. However, the computational gain of these MOM estimators over the likelihood estimators is significant for large *n* and *M* and often outweighs limitation of large standard error. There was no apparent bias in these estimators besides the repeat structure in Simulation 1C. For repeat structures of the causal markers, we observed underestimation in HE regression and a small upward bias for GCTA estimator.

## 5 Supplementary Material

Supplementary material is available online at http://biostatistics.oxfordjournals.org.

## 6 Data Availability

Derived data supporting the findings of this study are available on request. The code used to generate the data is available here: https://github.com/alantmin/heritability.

## 7 Acknowledgements

This work was supported in part by NIH/NIDA R21DA046188. Alan Min was additionally supported by the Statistical Genetics Training Grant T32 GM081062. This material is based upon work supported by the National Science Foundation Graduate Research Fellowship under Grant No. DGE-1762114. *Conflict of Interest*: None declared.

## Notes

### Competing Interest Statement

The authors have declared no competing interest.

