## Supplementary Materials for "Comparing Heritability Estimators under Alternative Structures of Linkage Disequilibrium"

### Supplementary material

#### S1 Likelihood-based Approaches:

The **GCTA REML** (Yang et al., 2011) estimator is derived by assuming that random SNP effects $\boldsymbol{\beta} \sim N(0, \sigma_g^2 \boldsymbol{I}_{m\times m})$ and that the normalized genotypes $\boldsymbol{\Gamma}$ are fixed. It assumes a random-effect model based approach for generation of phenotypes, and uses a Euclidean distance kernel for GRM calculation. Using the Normality assumption of $\boldsymbol{\beta}$, the GCTA REML estimator assumes that $y \sim N(0, \sigma_g^2\Psi + \sigma_e^2 I)$ and uses a restricted maximum likelihood (REML) approach to estimate $\sigma_g^2$ and $\sigma_e^2$. Recently, binning methods, such as in GCTA-LDMS have been used to apply GCTA on markers binned for different linkage disequilibrium (LD) structures or for different allele frequencies (Yang et al., 2015). However, such binning techniques are somewhat adhoc and are not incorporated in our simulation and analytical derivations.

The **LDAK** (Speed et al., 2012, 2017) estimator uses a similar approach to the GCTA REML estimator, also assuming fixed genotypes and random $\boldsymbol{\beta}$. The LDAK model tries to correct for uneven LD by computing a reweighted GRM as in Equation (S1).

$$X_{ij} = (G_{ij} - 2f_j) \times [2f_j(1 - f_j)]^{\alpha} \tag{S1}$$

The value $\alpha = -1.25$ is reported to generally work well with genomewide LD structure. Each of the raw genotypes is then weighted by substituting each column of $G_j$ with $w_j G_j$, where $w_j$ is chosen so that

$$w_j + \sum_j^{\prime} w_{j'} r_{jj'}^2 e^{-\lambda d_j j'} \tag{S2}$$

is constant over $j$. The squared correlation coefficient between SNPs $j$ and $j'$ is denoted by $r_{jj'}^2$, the genomic distance is denoted by $d_{jj'}$, and $\lambda$ is a constant. Note that $\alpha = -1$ corresponds with the GCTA REML estimator if all $w_j$ are 1.

#### S2: Method of Moments Estimators: no-LD

In this section we derive basic moment properties of the random-effect Haseman-Elston (HE) estimator and the fixed-effects Dicker-1 estimator in the case of no LD. We see their differences, but also their similarity in practice. The more general case with LD is considered in Section 2.3 of the main paper.

#### S2.1 Haseman Elston Method of Moments Estimator:

The HE estimator is a second-order moments estimator based on a regression of products of phenotypes $y_i y_k$ for all pairs $i \neq k$ on the corresponding $(i,k)$ terms of the $n \times n$ GRM matrix $\boldsymbol{\Psi} = M^{-1}\boldsymbol{\Gamma}_A \boldsymbol{\Gamma}'_A$. Given the standardized genotypes $\boldsymbol{\Gamma}$, the phenotypes depend only on the first $m$ causal markers and $\mathbf{y} = \boldsymbol{\Gamma}_C\ \boldsymbol{\beta} + \boldsymbol{\epsilon}$, where the independent variables $\beta_j \sim N(0, \sigma_g^2/m)$, and $\epsilon_i \sim N(0, \sigma_e^2)$.

We first consider the estimator as a regression estimate conditional on $\Psi$. Noting $i \neq k$, so $\mathrm{E}(\epsilon_\mathrm{i}\ \epsilon_\mathrm{k}) \ = \ 0$ and that the $\beta_j$ are independent, with mean 0 and variance $\sigma_g^2/m$,

$$\mathrm{E}(\mathrm{y_i y_k}\Psi_\mathrm{ik}) = \mathrm{E}\left( (\sum_{\mathrm{j=1}}^{\mathrm{m}} \Gamma_\mathrm{ij}\beta_\mathrm{j})\ (\sum_{\ell=1}^{\mathrm{m}} \Gamma_{\mathrm{k}\ell}\beta_\ell)\Psi_\mathrm{ik} \right) = \mathrm{E}\left( \sum_{\mathrm{j=1}}^{\mathrm{m}} \Gamma_\mathrm{kj}\Gamma_\mathrm{ij}\mathrm{E}(\beta_\mathrm{j}^2)\Psi_\mathrm{ik} \right) = \Psi_\mathrm{ik}^2\ \sigma_\mathrm{g}^2$$

Summing over all $n(n-1)/2$ pairs of distinct individuals, we have the method-of-moments equation

$$S_{Y\Psi} \quad \equiv \quad \sum_k \ \sum_{i<k} y_i y_k \Psi_{ik} \quad = \quad \sigma_g^2 \sum_k \ \sum_{i<k} \Psi_{ik}^2 \quad \equiv \quad \sigma_g^2 S_{\Psi\Psi}$$

so that $\sigma_g^2$ may be estimated as

$$\widetilde{\sigma_g^2} \ = \ \frac{S_{Y\Psi}}{S_{\Psi\Psi}} \ = \ \frac{\sum_k \sum_{i<k} y_i y_k \Psi_{ik}}{\sum_k \sum_{i<k} \Psi_{ik}^2} \tag{S3}$$

Then an estimate of heritability is given by dividing by the empirical variance of $\mathbf{y}$.

Here we focus on the estimate of $\sigma_g^2$ and on the numerator and denominator denoted $S_{Y\Psi}$ and $S_{\Psi\Psi}$ respectively. We consider not only the conditional model, but also the variation in $\Psi$ over samples of genotypes from the population. Note that

$$\Psi_{ik} = M^{-1} \sum_{j=1}^{M} \Gamma_{ij}\Gamma_{kj} \quad \text{and} \quad \mathrm{E}(\Gamma_\mathrm{ij}) = 0, \ \ \mathrm{E}(\Gamma_\mathrm{ij}^2) = 1$$

So if individuals are independent, $\mathrm{E}(\Psi_\mathrm{ik}) = 0$, and if markers are independent,

$$\mathrm{E}(\Psi_\mathrm{ik}^2) = \mathrm{var}(\Psi_\mathrm{ik}) = \mathrm{M}^{-1}\mathrm{var}(\Gamma_\mathrm{ij}\Gamma_\mathrm{kj}) \ = \ \mathrm{M}^{-1}(\mathrm{E}(\Gamma_\mathrm{ij}^2))^2 \ = \ 1/\mathrm{M}$$

and, under independence of individuals $i,k$ and independence of markers $j,w,\ell$,

$$\begin{aligned}
\mathrm{E(y_iy_k\Psi_{ik})} &= M^{-1}\ \mathrm{E}\left( (\sum_{\mathrm{j=1}}^{\mathrm{m}} \Gamma_{\mathrm{ij}}\beta_{\mathrm{j}} + \epsilon_{\mathrm{i}})(\sum_{\mathrm{w=1}}^{\mathrm{M}} \Gamma_{\mathrm{iw}}\Gamma_{\mathrm{kw}})(\sum_{\ell=1}^{\mathrm{m}} \Gamma_{\mathrm{k}\ell}\beta_\ell + \epsilon_{\mathrm{k}}) \right) \\
&= M^{-1}\mathrm{E}\left( \sum_{\mathrm{j=1}}^{\mathrm{m}} \beta_{\mathrm{j}}^2 (\sum_{\mathrm{w=1}}^{\mathrm{M}} \Gamma_{\mathrm{ij}}\Gamma_{\mathrm{iw}}\Gamma_{\mathrm{kw}}\Gamma_{\mathrm{kj}}) \right) \\
&= M^{-1}\mathrm{E}\left( \sum_{\mathrm{j=1}}^{\mathrm{m}} \beta_{\mathrm{j}}^2 \Gamma_{\mathrm{ij}}^2 \Gamma_{\mathrm{kj}}^2 \right) \quad = \quad \mathrm{M^{-1}m(\sigma_g^2/m) = \sigma_g^2/M}
\end{aligned}$$

Hence $S_{\Psi\Psi}$ has expectation $n(n-1)/2M$ and $S_{Y\Psi}$ has expectation $\sigma_g^2 n(n-1)/2M$. Empirical simulations (not shown) showed that while the standard deviation of $S_{\Psi\Psi}$ is approximately $n/M$, that of $S_{Y\Psi}$ is of order $n/\sqrt{M}$, but both decrease to 0 as $M \to \infty$. Thus as $M \to \infty$ with n remaining fixed, both $S_{Y\Psi}$ and $S_{\Psi\Psi}$ converge in probability to 0. As the number of markers increases, the coefficient of variation of $S_{\Psi\Psi}$ remains constant, but that of $S_{Y\Psi}$ increases, and the empirical study shows the the standard deviation of the estimate of $\sigma_g^2$ to be of order $\sqrt{M}/n$. This result is in agreement with the theoretical equations for the estimator of Dicker (2014) in the case of no LD: see Lemma 2 and the Remarks following in that paper. That is, uncertainty in $\sigma_g^2$ and hence in $h^2$ increases as the number of markers $M$ increases.

#### S2.2 The Dicker-1 fixed-effects model moments estimator

The Dicker-1 estimator (Dicker, 2014) is also a method of moments estimator, but starts from very different assumptions. The standardized genotypes $\Gamma_{ij}$ are are assumed to be distributed $N(0,1)$, independent over individuals $i$. The effects $\beta_j$ are fixed effects, and in our case where only the first $m$ markers are causal, $\beta_j \equiv 0$ for $j = (m+1), ...M$. The parameter to be estimated is $\sigma_g^2 \equiv \boldsymbol{\beta}'\boldsymbol{\Sigma}\boldsymbol{\beta}$ where here $\boldsymbol{\beta}$ is the $m$-vector of effects at causal markers augmented by $(M-m)$ zeros and $\boldsymbol{\Sigma}$ is the LD matrix of correlations among all $M$ markers. Because of the Normality assumption for genotypes, these can be rotated to orthonormality. This implies that the case of known $\boldsymbol{\Sigma}$ is mathematically equivalent to $\boldsymbol{\Sigma} = \boldsymbol{I}$. For simplicity we consider this case, then $\boldsymbol{\beta}'\boldsymbol{\Sigma}\boldsymbol{\beta} = \sum_{j=1}^m \beta_j^2$ and $m^{-1}\sum_{j=1}^m \beta_j^2 \ \equiv\ \sigma_g^2/m$, equivalent, for large $m$ to the random-effects HE assumption $\beta_j \ \sim\ N(0, \sigma_g^2/m)$.

Dicker (2014) uses the quadratic forms $\|\mathbf{y}\|^2 = \mathbf{y'y}$ and $\|\mathbf{\Gamma}'_A\mathbf{y}\|^2 = M\ \mathbf{y'\Psi y}$. Without making Normality assumptions, we can compute

$$\begin{aligned}\mathrm{E}(\mathrm{M}\ \mathbf{y'\Psi y}) = & \sum_{i=1}^{n}\sum_{k=1}^{n}\mathrm{E}(\mathrm{M\ y_i\Psi_{ik}y_k}) \\ = & \sum_{i=1}^{n}\sum_{k=1}^{n}\mathrm{E}\left((\sum_{\mathrm{j=1}}^{\mathrm{m}}\Gamma_{\mathrm{ij}}\beta_{\mathrm{j}}+\epsilon_{\mathrm{i}})(\sum_{\mathrm{w=1}}^{\mathrm{M}}\Gamma_{\mathrm{iw}}\Gamma_{\mathrm{kw}})(\sum_{\ell=1}^{\mathrm{m}}\Gamma_{\mathrm{k}\ell}\beta_{\ell}+\epsilon_{\mathrm{k}})\right)\end{aligned} \tag{S4}$$

Under independence of $\Gamma_{iw}$ and $\Gamma_{kw}$ for $i \neq k$, the coefficient of $\sigma_e^2$ is seen to be $Mn$. Under independence of markers indexed by $j$, $\ell$ and $w$, the majority of terms in $\beta_j$ and $\beta_l$ in this expression disappear, leaving only a coefficient of $\sigma_g^2 = \sum_{j=1}^m \beta_j^2$. The remaining terms have $j = \ell \neq w$ (in which case $i = k$), or $j = \ell = w$ (in which case terms with both $i = k$ and $i \neq k$ remain). Grouping these two sets of terms this coefficient reduces to

$$(M-1)\ \mathrm{E}(\sum_{\mathrm{i}}\Gamma_{\mathrm{ij}}^2\Gamma_{\mathrm{iw}}^2)+\mathrm{E}(\sum_{\mathrm{i}}\sum_{\mathrm{k}}\Gamma_{\mathrm{ij}}^2\Gamma_{\mathrm{kj}}^2) \quad = \quad \mathrm{n(M-1)+Kn+n(n-1)} \ = \ \mathrm{n(M+n+K-2)}$$

where $K = \mathrm{E}(\Gamma_{\mathrm{ij}}^4)$. Combining the following two equations,

$$\begin{aligned}\mathrm{E}(\mathrm{n}^{-1}\mathrm{M}\ \mathbf{y'\Psi y}) &= (M+n+K-2)\sigma_g^2\ +\ M\sigma_e^2 \\ \mathrm{E}(\mathrm{n}^{-1}\mathbf{y'y}) &= \sigma_g^2+\sigma_e^2\end{aligned}$$

and assuming $K = 3$ we obtain the Dicker (2014) method-of-moments estimator of $\sigma_g^2$:

$$\tilde{\sigma}_g^2 \ = \ (n(n+1))^{-1}(M\ \mathbf{y'\Psi y}\ -\ M\mathbf{y'y}) \ = \ (n(n+1))^{-1}(\|\mathbf{\Gamma}'_A\mathbf{y}\|^2\ -\ M\|\mathbf{y}\|^2) \tag{S5}$$

Note that whereas the numerator and denominator of the HE estimator (6) always has the correct expectations, Equation (S5) is only exact if $K = 3$. Since $K$ appears only in the term $(M + n + K - 2)$ the impact will be small for large $M$ and/or $n$, but it is worth noting that $K$ can be quite large ($> 100$) for loci with rare alleles (see Figure S1). Under the $N(0, 1)$ assumption, Dicker (2014) gives also many other expressions for high-order moments of these estimators. However, these depend more critically on the higher-order moments of the $\Gamma_{ij}$, and hence his Normality assumption.

Although the assumptions underlying the MoM estimator (S5) are very different from those of the HE estimator of Equation (6), operationally and in performance the estimators are quite similar, in the case of

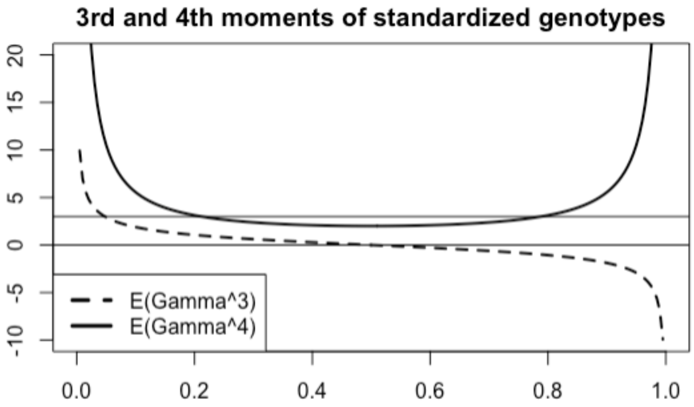


Figure S1: Skewness and kurtosis of the normalized genotypes as a function of allele frequency

known or no LD. The key difference from the HE estimator is then that whereas the latter considers only $\Psi_{ik}$ for $i \neq k$, the Dicker estimator uses the full $n \times n$ matrix $M\mathbf{\Psi} = \mathbf{\Gamma}_A\mathbf{\Gamma}'_A$. This use of the diagonal terms $\Psi_{ii}$ permits an estimators of $\sigma_g^2$ and $\sigma_e^2$ that is linear in the relevant quadratic forms, rather than the ratio $S_{YT}/S_{TT}$, but strict correctness and moment properties are dependent on the Normality assumption for $\Gamma_{ij}$.

### S3: Moment based estimators: LD case

#### S3.1 Biases in HE estimator in the presence of LD

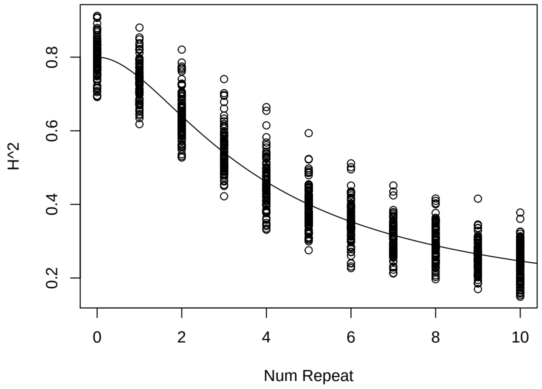


Figure S2: Estimates of $h^2$ using the HE estimator are shown with a true heritability of 0.8 on the Y-axis. The solid line plots the theoretical estimates based on Equation (14). 1000 individuals and 200 markers were simulated, where the markers were simulated from the binomial distribution using allele frequencies from the AFR population from 1000 genomes. 20 markers were repeated for “Num Repeat” times, as plotted on the X-axis.

In the presence of LD the HE estimator may be biased. A formula for this bias, approximating the expectation of a ratio by the ratio of expectations, is derived in Section 2.3 of the main paper. Of the simulation study examples of this paper, the bias is marked in the case of non-causal markers that repeat the genotypes of causal markers. Figure S2 aims to validate the formula for the bias and assess the variation in bias across realizations.

#### S3.2 Moment estimators designed to accommodate LD

In the case when LD must be estimated from the sample data, Dicker (2014) and Schwartzman et al. (2019) developed moment-based estimators of $\sigma_g^2$, $\sigma_e^2$, and $h^2$ under the fixed-effects framework.

Here we consider the estimator of Dicker (2014) in the case of LD. Again, the GRM $\Psi = M^{-1}\mathbf{\Gamma}_A\,\mathbf{\Gamma}'_A$, and LD matrix $\mathbf{\Sigma} = n^{-1}\mathbf{\Gamma}'_A\mathbf{\Gamma}_A$. If the standardized genotypes, $\Gamma_{ij}$, are marginally $N(0,1)$ and independent over $i$, and if $\mathbf{\Sigma}$ is the true positive definite correlation matrix of the $\Gamma_{ij}$ over $j$, then $\mathbf{\Sigma}^{-1/2}\mathbf{\Gamma}'_A$ are independent $N(0,1)$ and the estimator (S5) becomes

$$\begin{aligned}\tilde{\sigma}_g^2 &= (n(n+1))^{-1}((\mathbf{\Sigma}^{-1/2}\mathbf{\Gamma}'_A\mathbf{y})'(\mathbf{\Sigma}^{-1/2}\mathbf{\Gamma}'_A\mathbf{y}) \;-\; M\mathbf{y}'\mathbf{y}) \\ &= (n(n+1))^{-1}(\mathbf{y}'\mathbf{\Gamma}_A\mathbf{\Sigma}^{-1}\mathbf{\Gamma}'_A\mathbf{y} \;-\; M\mathbf{y}'\mathbf{y}) \qquad \text{(S6)}\end{aligned}$$

and again $\sigma_g^2 + \sigma_e^2$ is estimated by the phenotypic variance $n^{-1}\mathbf{y}'\mathbf{y}$. More generally, as shown by Dicker (2014), if $n > M$ and $\mathbf{\Sigma}$ is a norm-consistent estimator of the true correlation matrix the properties and results of the non-LD estimator (S5) apply also in the LD case to the estimator (S6).

However, in most applications, $M$ is much larger than $n$. and the estimator (S6) breaks down, and as shown in Dicker (2014), In this case they propose to use lower-order moments of the trace of $\mathbf{\Sigma} \;=\; n^{-1}\mathbf{\Gamma}'_A\mathbf{\Gamma}_A$. Specifically they define

$$\mu_1 \;=\; \frac{tr(\mathbf{\Sigma})}{M} \text{ and } \mu_2 \;=\; \frac{tr(\mathbf{\Sigma}^2)}{M} - \frac{(tr(\mathbf{\Sigma}))^2}{Mn} \qquad \text{(S7)}$$

The estimator of $\sigma_g^2$ becomes

$$\tilde{\sigma}_g^2 \;=\; \frac{\mu_1(\mathbf{\Gamma}'_A\mathbf{y})'(\mathbf{\Gamma}'_A\mathbf{y}) \;-\; M\mu_1^2\mathbf{y}'\mathbf{y}}{n(n+1)\mu_2} \qquad \text{(S8)}$$

and again $\sigma_g^2 + \sigma_e^2$ is estimated by $n^{-1}\mathbf{y}'\mathbf{y}$. For more on the theory and properties of the estimator (S8) see Dicker (2014). For the current paper, we implement this estimator as “Dicker-2” in our simulations and results.

Schwartzman et al. (2019) proposed a method of moments estimator based on that of Dicker (2014).

They derive a form that depends only on summary statistics instead of the raw genotypic and phenotypic data and hence their estimator has wider applicability. However, in the basic form (not using only summary statistics) their estimator is essentially equivalent to the estimator (S8), so we do not consider it further in this paper.

#### S4. Simulation of Genetic Marker LD Structures

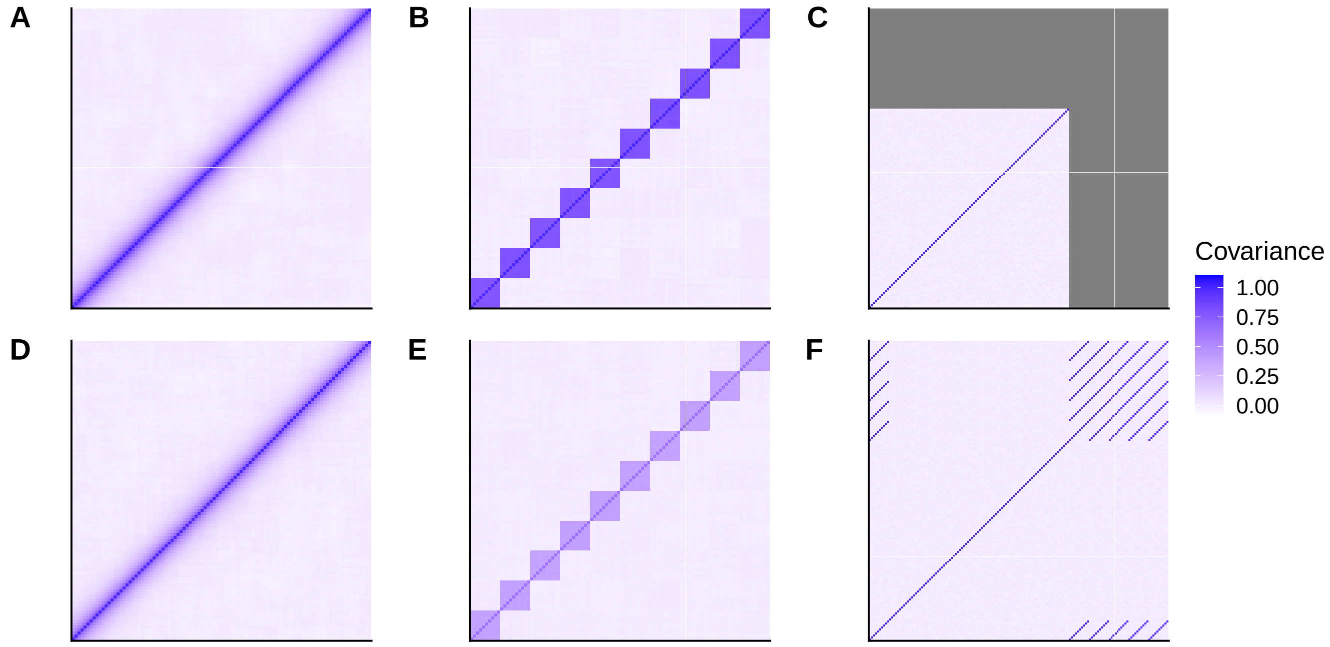


Figure S3: These panels plot the empirical covariance matrices for simulated genotypes from 10,000 individuals and $p = 100$ markers. The correlation between markers decreases after discretization but the pattern generally remains the same. (A) Autocorrelated markers were generated from the Gaussian model, i.e. plotting $Cov(\tilde{G})$ (B) Blocked markers were generated from the Gaussian model. (C) Independent markers were generated. (D) Autocorrelated markers were generated and then discretized and normalized, i.e. this is $Cov(\Gamma)$ (E) Blocked markers were discretized and normalized. (F) Repeated markers were generated with 10 markers being repeated 5 times.

**Autocorrelated:** we assume that for each individual, $M$ markers are generated from a multivariate Gaussian with $AR1(\rho)$ covariance matrix. We generate the markers for each individual independently. In other words, we assume that for individual $i$, genotypes $\tilde{G}_i$ are generated from $\tilde{G}_i \sim N(0, \Sigma)$, where

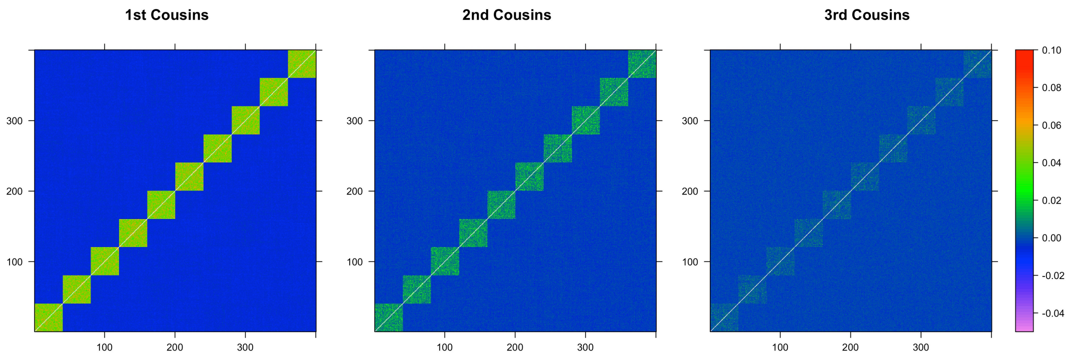


Figure S4: Colors represent values of the log of 1 plus the average of 100 GRMs generated from 400 individuals. The $i, j$th entry of the matrix corresponds to the relatedness between $i$th individual and the $j$th individual. Sets of cousins are adjacent in groups of 40. Colors are thresholded at 0.1, and set to white if it is above the threshold.

$$\Sigma = \begin{pmatrix} 1 & \rho & \rho^2 & \dots & \rho^{M-1} \\ \rho & 1 & \rho & \dots & \rho^{M-2} \\ \vdots & & \vdots & & \vdots \\ \rho^{M-1} & \rho^{M-2} & \rho^{M-3} & \dots & 1 \end{pmatrix}$$

The continuous values $\tilde{G}_i$ are then converted to discrete genotypes $G_i$ taking value 0, 1 or 2. For a marker with alternate allele frequency $f$, $G_{ij} = 0$, 1, or 2, depending on if $\tilde{G}_{ij}$ is less than $\Phi^{-1}(f^2)$, between $\Phi^{-1}(f^2)$ and $\Phi^{-1}(f^2 + 2f(1-f)) = \Phi^{-1}(2f - f^2)$, or greater than $\Phi^{-1}(2f - f^2)$, where $\Phi(\cdot)$ is the $N(0,1)$ distribution function. Note that this trichotomy gives the correct marginal genotype probabilities, but reduces the genotypic correlation (LD) between markers below that used in the simulation matrix $\Sigma$: compare panels A with D, or B with E in Figure S3.

**Block:** we generate block genotypes according to the same mechanism as the autocorrelated genotypes, except we choose that

$$\Sigma = \begin{pmatrix} 1 & \rho & \rho & \dots & \rho \\ \rho & 1 & \rho & \dots & \rho \\ \vdots & & \vdots & & \vdots \\ \rho & \rho & \rho & \dots & 1 \end{pmatrix}$$

for each block. We assume that there are 10 blocks, each with $M/10$ markers.

**Repeat:** In this case $m$ marker genotypes are independently generated from the binomial distribution.

That is, for a marker with alternate allele frequency $f$, $G_{ij} \sim Binomial(2, f)$. We designate a proportion of markers to be repeated. We repeat these markers $r$ times.

**Choice of causal markers**

For the three simulation LD structures, we selected $\boldsymbol{G}_c$ to be a subset of $\boldsymbol{G}$. For the autocorrelation and block simulated genotypes, we chose alternating markers to be causal and non-causal markers. For the repeat structure the original $m$ markers were chosen to be causal, while the repeat genotypes were non-causal. The genotypes were standardized to each have mean 0 and variance 1, using the empirical allele frequencies in the simulated sample of $n$ individuals. The matrix $\mathbf{\Gamma}_A$ of standardized genotypes was formed as given in Equation (1), while $\mathbf{\Gamma}_C$ is the corresponding matrix for the $m$ causal markers.
